# Sphingosine kinase 2 suppresses neutrophil responses to promote viral persistence while attenuating immune pathology

**DOI:** 10.1101/2025.09.08.674905

**Authors:** Vijayamahantesh Vijayamahantesh, Ying He, Lei Jiang, Hailey Huerter, Kwang Il Jung, Savannah McKenna, Caleb J. Studstill, Lee-Ann H. Allen, Ravi Nistala, Dong Xu, Bumsuk Hahm

**Affiliations:** Departments of Surgery and Molecular Microbiology & Immunology, University of Missouri, Columbia, MO 65212, USA; Department of Electrical Engineering and Computer Science, and Christopher S. Bond Life Sciences Center, University of Missouri, Columbia, MO 65211, USA; Department of Molecular Microbiology & Immunology, University of Missouri, Columbia, MO 65212, USA; Division of Nephrology, Department of Medicine, University of Missouri, Columbia, MO 65201, USA

## Abstract

Chronic virus infections often suppress immune cell functions which helps in restricting immune pathology but leads to viral persistence. However, the underlying mechanisms are incompletely understood. We recently found that sphingosine kinase 2 (SphK2)-deficient (*Sphk2*^-/-^) mice succumbed to lymphocytic choriomeningitis virus (LCMV) infection due to immune pathology. In addition to heightened T cell immunity, a notable increase of neutrophils was observed in LCMV-infected *Sphk2*^-/-^ mice. Depletion of neutrophils increased the viability of virus-infected *Sphk2^-/-^* mice, supporting a role of SphK2-deficient neutrophils in viral immune pathogenesis. Further, SphK2-deficient neutrophils expressed lower levels of the immune suppressive marker CD244 during infection. Importantly, adoptively transferred SphK2-deficient neutrophils demonstrated intrinsic regulation of CD244 and improved virus-specific T cell responses, resulting in diminished viral burden. Transcriptomic analysis revealed increased expression of pro-inflammatory and antiviral genes in SphK2-deficient neutrophils. These results indicate that SphK2 promotes suppressive neutrophil responses and regulates neutrophil-associated immune pathology during a persistent infection. Our findings may help design new immune therapeutics to control chronic viral diseases.

**Significance:** Neutrophils are the sentinels of the innate immune system; they can reshape innate and adaptive immune responses. During chronic illnesses, such as persistent viral infections, neutrophils can suppress the host immune response and help in disease progression. Here, we demonstrate regulation of neutrophil expansion and functions by sphingosine kinase 2 (SphK2) during LCMV infection. SphK2-deficient neutrophils express a reduced level of inhibitory receptor CD244, exert immune stimulatory effects on T cells, and promote virus clearance. Further, transcriptomic analysis reveals that SphK2 deficiency leads to the development of proinflammatory neutrophils. Our study identifies SphK2, a host factor, as being critical for neutrophil suppression that regulates dysfunctional T cell response and virus persistence.

## Introduction

Viruses can establish persistent infections, causing chronic and fatal illnesses that account for almost a million deaths per year (1). Chronic viral infections often induce immune suppressive conditions by regulating diverse immune cells and factors. Viral immune suppression is exemplified by the gradual loss of effector T cell functions (2). However, the molecular and cellular mechanism of immune dysfunction during chronic infections is incompletely understood.

Lymphocytic choriomeningitis virus (LCMV) infection of mice has been useful for studying the host immune response to infections and for understanding key immunological mechanisms (3–9). LCMV prototypic strain Armstrong (Arm) causes acute infection, as the virus elicits a potent CD8^+^ effector T cell response that results in subsequent virus clearance within a week. On the other hand, LCMV clone 13 (Cl 13), a variant of the parental Arm strain, induces immune suppression, which allows the virus to persist up to 60-100 days post-infection (10–13). Mouse models using LCMV Cl 13 have also been instrumental in uncovering immunological concepts such as virus-specific T cell exhaustion (12–15). These concepts held true during other chronic infections and diseases in humans and significantly contributed to the development of the first immunotherapy targeting PD-1 for the treatment of cancers (16).

Sphingolipids are bioactive lipid molecules, as they are not only essential structural components of the lipid bilayer, but also key players in intra- and extracellular signaling systems (17). Sphingosine 1-phosphate (S1P) is a sphingolipid metabolite and an important regulator of inflammation and immune responses (18, 19). Generation of S1P from sphingosine is catalyzed by sphingosine kinase 1 (SphK1) and sphingosine kinase 2 (SphK2). These two enzymes are encoded by two genes and differ in their subcellular localization, substrate specificity, and roles. SphK1 is mainly localized in the cytosol and translocates to the plasma membrane upon activation (20). S1P produced by SphK1 on the plasma membrane acts in an autocrine or paracrine manner to promote cell survival and proliferation. Although SphK2 is found in the cytoplasm, it shuttles between the nucleus and other subcellular organelles and is known to regulate epigenetic gene expression (21). Despite sharing enzymatic function, SphK1 and SphK2 differentially regulate immune responses through mechanisms not well defined (22).

Previously, we have shown that deletion of SphK2 increased T cell response, immune pathogenesis, and eventually death of LCMV Cl 13-infected mice due to vascular leakage in the kidney (23). Of note, we also observed a significantly increased neutrophil population in the kidney of LCMV Cl 13-infected SphK2-deficient mice compared to their wild-type (WT) counterparts. Historically, neutrophils have been seen as sentinels of the immune system and the first cells to reach the site of infection or inflammation. However, recent studies demonstrated that neutrophils could function in diverse ways, exhibit transcriptional and functional plasticity in response to extracellular cues, and participate in the shaping of innate and adaptive immune responses (24). Myeloid cells such as myeloid-derived suppressor cells (MDSCs) were reported to play a key role in T cell exhaustion, hence the development of virus persistence (25–27). However, the mechanism of immune suppression by myeloid cells, especially neutrophils, is not clear.

Here, we have investigated the role of neutrophils during chronic LCMV infection. Our results demonstrate that SphK2 intrinsically regulates the expression of immune regulatory molecule CD244 on neutrophils during infection, and SphK2-deficient neutrophils exhibited immune stimulatory effects on T cells. Single-cell RNA-sequencing (scRNA-seq) of bone marrow neutrophils (BMNs) revealed the role of SphK2 in the regulation of several key genes required for the development of immune suppressive neutrophils. Taken together, these results suggest SphK2 can be a valuable therapeutic target against chronic infections or inflammatory conditions.

## Results

### SphK2 deficiency induces expansion of neutrophils upon LCMV Cl 13 infection which contributes to viral immune pathogenesis

LCMV Cl 13 infection of SphK2-deficient mice resulted in increased T cell responses associated with immune pathology in kidneys and eventually led to the death of infected mice. While SphK2 deficiency scarcely affected other immune cell components, we observed increased infiltration of neutrophils into the kidneys of *Sphk2^-/-^* mice compared to their wild-type (WT) mice upon infection(23). As neutrophils are often responsible for inflammation and tissue damage, the neutrophil responses in *Sphk2^-/-^* mice were further assessed during Cl 13 infection. The percentage and distribution of neutrophils can quickly change during infection, affecting the establishment of inflammation and immune responses (28). We assessed the changes in neutrophil population during chronic LCMV infection. As shown in Fig. 1, neutrophils in spleen, blood, and kidney of *Sphk2^-/-^* mice significantly increased compared to WT mice at 3 days post-infection (dpi) (Fig. 1A) and 8 dpi (Fig. 1B). A neutrophil surge in spleen, blood, and kidney of *Sphk2^-/-^* mice at 3 dpi was in line with the significantly increased neutrophils in BM of *Sphk2^-/-^*mice (Fig. 1A and 1C). Interestingly, at 8 dpi, although there was no significant difference in neutrophils in the primary lymphoid organ BM, we still observed significantly increased neutrophils in the blood, spleen, and kidney (Fig. 1B). The increase in neutrophils in spleen and blood was sustained when assessed at 14 dpi (Supplementary Fig. S1). These results indicate that LCMV Cl 13 induces the early expansion of neutrophils in the *Sphk2^-/-^*mice, and this increased number of neutrophils persists during infection.

**Fig. 1.**
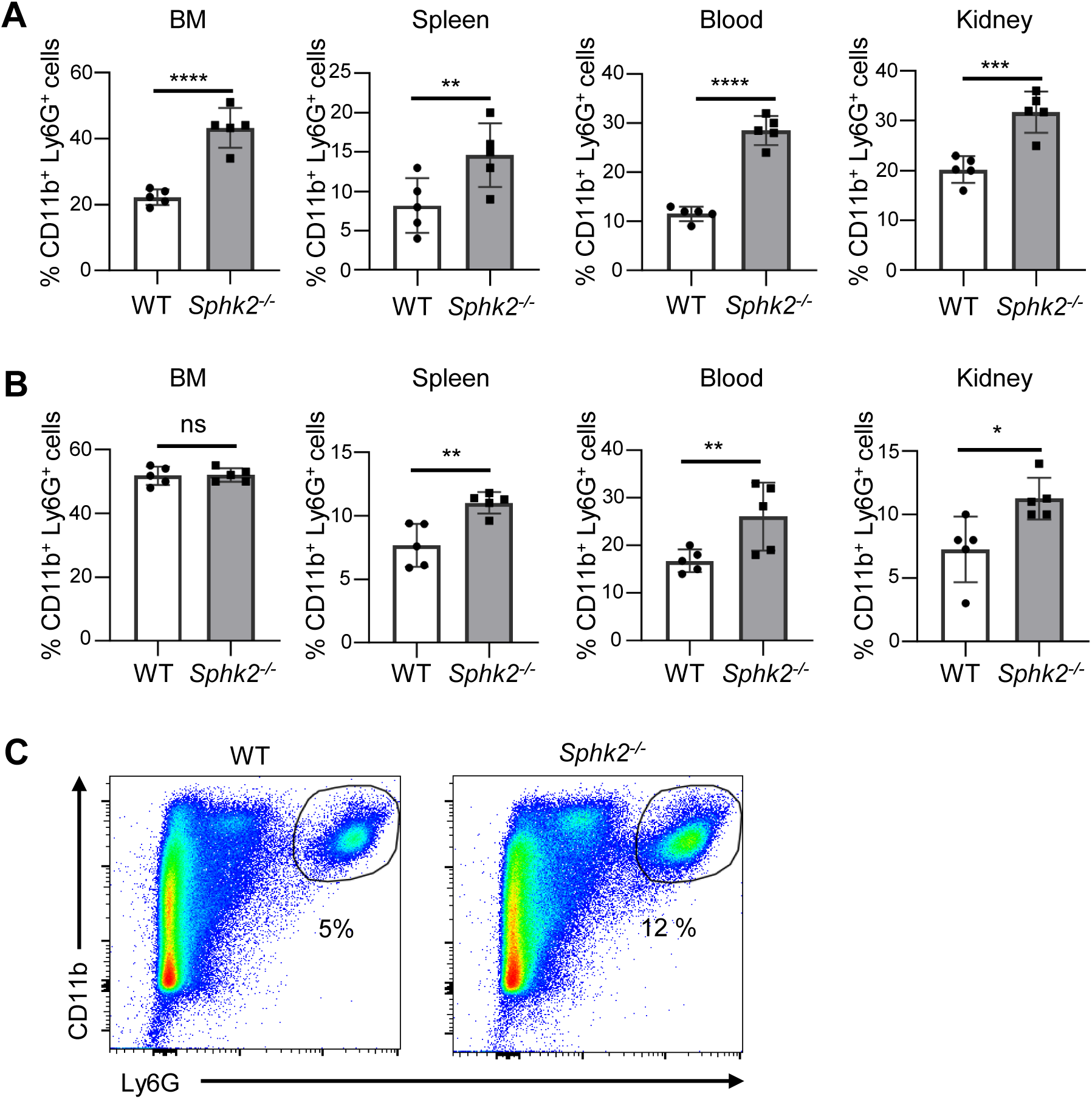
**Deletion of *Sphk2* induces an increase in neutrophils upon LCMV Cl 13 infection.** WT and *Sphk2^-/-^* mice (n = 5/group) were infected with LCMV Cl 13. Mice were sacrificed at 3 and 8 dpi, and organs were collected. The percentage of neutrophils (CD11b^+^Ly6G^+^) in bone marrow (BM), spleen, blood, and kidney of LCMV Cl 13 infected WT and *Sphk2^-/-^* mice at 3 (**A**) and 8 (**B**) dpi was quantified using flow cytometry. A representative flow cytometry data of splenic neutrophils from C57BL/6 WT and *Sphk2^-/-^* mice at 3 dpi is shown (**C**). ****p≤0.0001, ***p≤0.001, **p≤0.01, *p≤0.05, *n.s.* not significant, bidirectional, unpaired Student’s *t*-test. Data are representative of 2-3 independent experiments.

To determine if the increased neutrophil expansion observed in LCMV Cl 13-infected *Sphk2^-/-^* mice is an LCMV Cl 13 strain-specific event, *Sphk2^-/-^* and WT mice were infected with LCMV Arm or Cl 13. On day 10 post-infection, the neutrophil population in the spleen of infected mice was analyzed. We found a significantly increased number of neutrophils in the *Sphk2^-/-^* mice compared to WT mice, irrespective of strains used (Fig. 2A, 2B, and 2C). These results suggest that neutrophil expansion is regulated by SphK2 regardless of LCMV strain used. However, LCMV Cl 13 infection induced accumulation of a higher percentage of neutrophils than LCMV Arm infection (Supplementary Fig. S2A and S2B), implying that the neutrophils may play a role in immune regulation during Cl 13 infection.

**Fig. 2.**
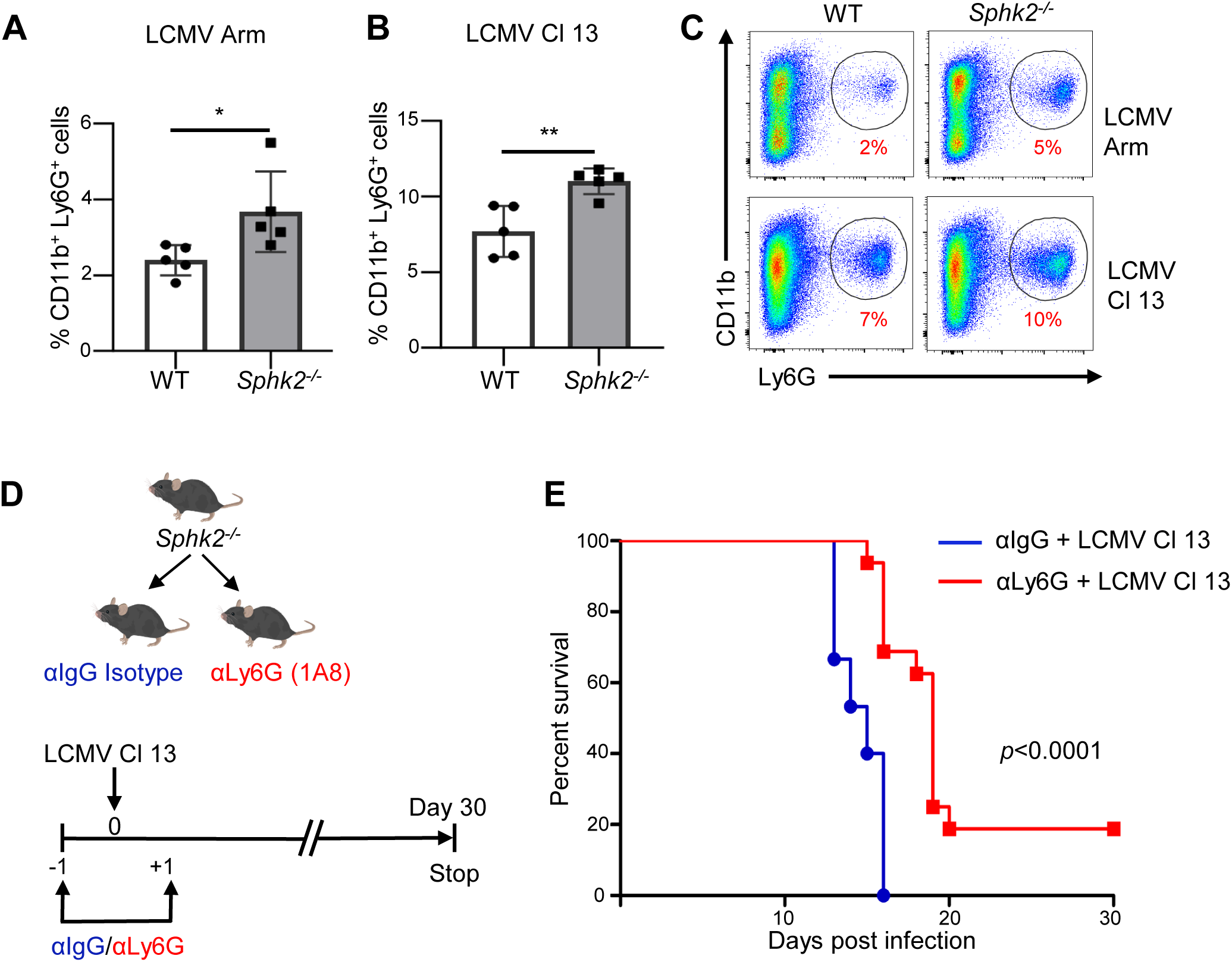
Increased neutrophil expansion by SphK2 deficiency is independent of LCMV strains and contributes to Cl 13-induced mortality of *Sphk2*^-/-^ mice. WT and *Sphk2^-/-^* mice (n = 5/group) were infected with LCMV Arm or Cl 13. At 10 dpi, the percentage of neutrophils (CD11b^+^Ly6G^+^) in the spleen of mice infected with LCMV Arm (**A**) and Cl 13 (**B**) was quantified by flow cytometry. A representative flow cytometry data of splenic neutrophils from WT and *Sphk2^-/-^* mice at 10 dpi is shown (**C**). Depletion of neutrophils was performed by injecting *Sphk2^-/-^* mice with the 250 µg anti-Ly6G antibody (αLy6G) or control-antibody (αIgG) via the intraperitoneal (i.p.) route one day before and one day after LCMV Cl 13 infection. The flow diagram of the neutrophil depletion strategy is shown in (**D**). The comparison of survival of isotype antibody and anti-Ly6G treated *Sphk2^-/-^* mice upon LCMV Cl 13 infection was monitored for 30 days (**E**). **p≤0.01, *p≤0.05, bidirectional, unpaired Student’s *t*-test for A and B and Kaplan-Meier test for E.

The systemic increase of neutrophils observed in LCMV Cl 13-infected *Sphk2^-/-^* mice led us to hypothesize that neutrophils contribute to the immune pathology of kidney damage and the ultimate mortality of LCMV Cl 13-infected *Sphk2^-/-^* mice. To test this, we adopted a neutrophil depletion strategy during LCMV Cl 13 infection. Using anti-Ly6G (1A8) antibody (αLy6G), neutrophils were depleted one day before and one day after LCMV Cl 13 infection followed by monitoring of mouse mortality, as shown in Fig. 2D. Depletion of neutrophils significantly increased survival of Cl 13-infected *Sphk2^-/-^* mice compared to the isotype control antibody (αIgG) treated group (Fig. 2E). The failure of complete rescue of LCMV Cl 13-infected *Sphk2^-/-^* mice by neutrophil depletion suggests the potential involvement of other immune cells, such as T cells, which were shown to be critical for virus-induced immune pathology(23). Taken together, these results indicate that SphK2 deficiency induces the expansion of neutrophils which contribute to immune pathogenesis.

### SphK2 intrinsically regulates CD244 expression on neutrophils upon LCMV Cl 13 infection

Although the lack of specific surface markers makes it challenging to separate neutrophil subpopulations, a few studies have exploited CD244 to differentiate suppressive neutrophils from resting or immune-stimulatory neutrophils (29–31). As a member of the signaling lymphocyte activation molecule family (SLAMF) receptors, CD244 can transmit an inhibitory or activation signal in lymphocytes, but generally transmit an inhibitory signal in myeloid cells due to the differential downstream regulatory pathways (32). Following LCMV Cl 13 infection, CD244 was upregulated on the surface of neutrophils in BM, spleen, blood, and kidney when compared to the uninfected group (Supplementary Fig. S3). To determine if SphK2 affects the expression level of CD244 on neutrophils during Cl 13 infection, CD244 expression on neutrophils (CD244^+^CD11b^+^Ly6G^+^ cells) was assessed in multiple organs from WT vs. *Sphk2^-/-^* mice at 3 dpi. The expression of CD244 on neutrophils and percentage of CD244^+^ neutrophils significantly decreased in BM, spleen, blood, and kidney in *Sphk2^-/-^* when compared to WT mice (Supplementary Fig.S4) (Fig. 3A). The representative plots show a clear difference in CD244^+^Ly6G^+^ cells between WT vs. *Sphk2*^-/-^ (75% vs. 27%) (Fig. 3B). These results indicate that SphK2 induces an increase of CD244 expression on neutrophils upon LCMV Cl 13 infection.

**Fig. 3.**
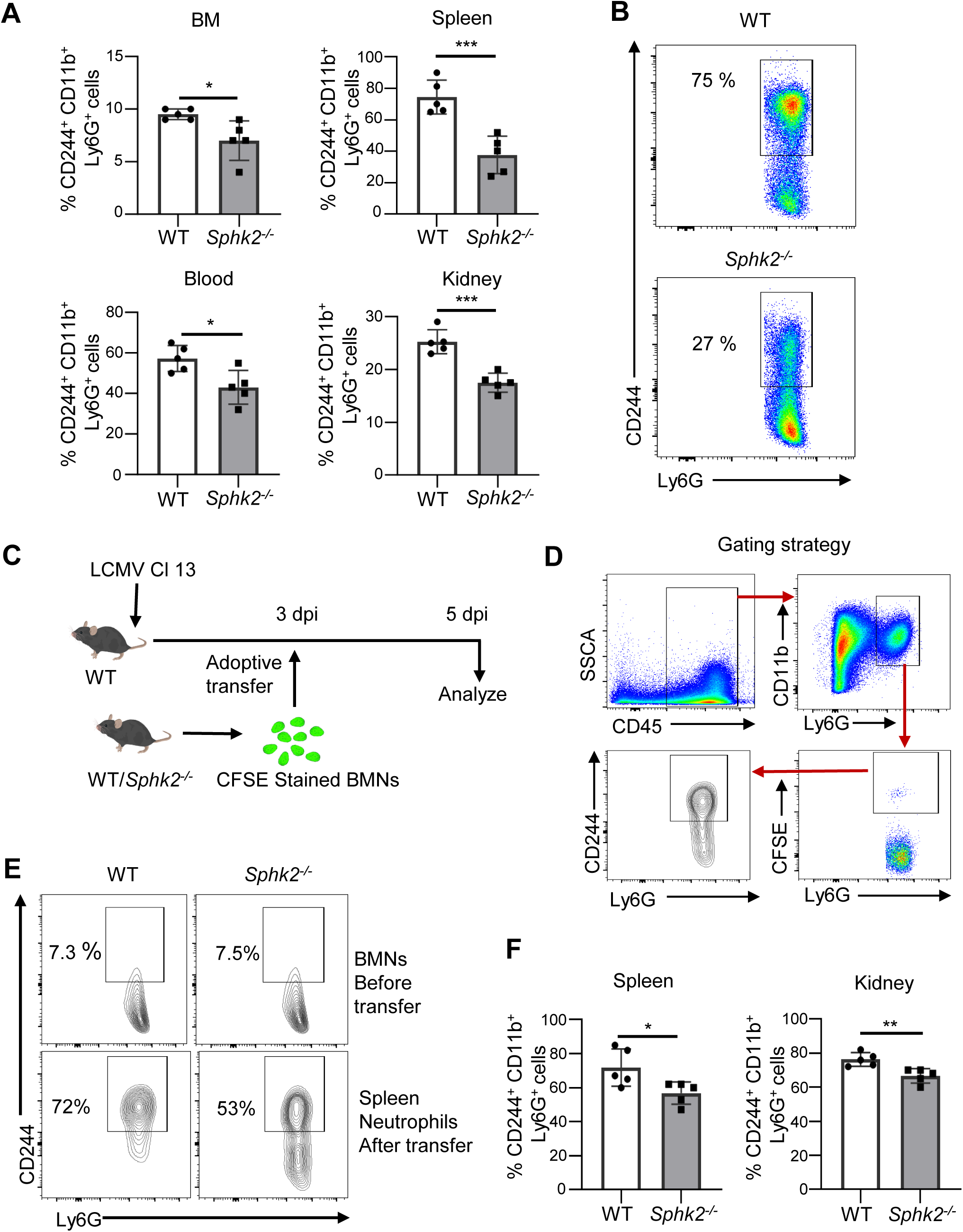
**Expression of CD244 is significantly reduced on *Sphk2^-/-^* neutrophils and is intrinsically regulated by SphK2**. WT and *Sphk2*^-/-^ mice (n = 5/group) were infected with LCMV Cl 13, and at 3 dpi, CD244^+^ neutrophils in organs were measured by flow cytometry (**A**). A representative flow cytometry data of CD244^+^ neutrophil between WT and *Sphk2^-/-^* mice on 3 dpi is shown (**B**). To determine the intrinsic regulation of CD244 expression by SphK2 on neutrophils, the experiments were performed by using adoptive transfer of donor BMNs (**C-F**). Naïve BMNs were isolated from WT and *Sphk2*^-/-^ mice (n = 5/group), stained with CFSE, and then adoptively transferred to LCMV Cl 13-infected WT mice at 3 dpi. The transferred cells were detected in the spleen and kidney at 5 dpi by flow cytometry. The flow diagram of neutrophil adoptive transfer (**C**) and the gating strategy of detecting CFSE-stained donor neutrophils in recipient mice (**D**) are depicted. Representative CD244^+^ neutrophil in donor BMNs before transfer (top) and on donor cells in the recipient spleen after transfer (bottom) are shown (**E**). The percentage of CD244 expression on CFSE-stained donor cells in the spleen and kidney of recipient mice was assessed (**F**). ***p≤0.001, **p≤0.01, *p≤0.05, bidirectional, unpaired Student’s *t*-test.

To investigate whether SphK2 can regulate CD244 expression in a neutrophil-intrinsic manner under LCMV infection condition, CFSE-stained BM neutrophils (BMN) from uninfected WT or *Sphk2*^-/-^ mice were adoptively transferred into LCMV Cl 13-infected WT mice on day 3 post-infection (Fig. 3C). The expression of CD244 on donor neutrophils (CFSE stained) in the spleen and kidney of recipient mice was assessed (Fig. 3C and 3D). Prior to adoptive transfer, the basal level of CD244^+^ neutrophils in donor BMNs was approximately 7% comparable between WT and *Sphk2*^-/-^ mice (Fig. 3E). However, when these neutrophils were exposed to the LCMV Cl 13 infection environment, the transferred WT neutrophils acquired a high level of CD244 expression. Consistent with the *Sphk2^-/-^* phenotype, SphK2-deficient neutrophils in the infection microenvironment showed reduced CD244^+^ neutrophils compared to those from WT donor mice (72% vs. 53% as shown in Fig. 3E and Fig. 3F). These observations indicate that SphK2 can regulate the expression of CD244 intrinsically on neutrophils during LCMV infection.

### *Sphk2^-/-^* neutrophils produce significantly higher ROS upon LCMV Cl 13 infection

Reactive oxygen species (ROS) production by neutrophils is one of the mechanisms of tissue damage that contributes to disease severity (33). Therefore, we determined if SphK2 deficiency alters the capacity of neutrophils to produce ROS by comparing the levels of ROS from BMNs of *Sphk2^-/-^* and WT mice following LCMV infection. ROS production was quantified by flow cytometry using the intracellular ROS detection probe H2DCFDA. Without LCMV infection, there was no significant difference at the basal level of ROS between WT and SphK2-deficient neutrophils in the presence or absence of PMA stimulation (Supplementary Fig. S5A and S5B). However, upon LCMV infection, we found significantly increased production of ROS by unstimulated neutrophils (Fig. 4A and 4B) as well as PMA stimulated neutrophils (Fig. 4C and 4D) from *Sphk2^-/-^* mice compared to WT mice. ROS production was also quantified in a time-dependent manner using a luminol-enhanced chemiluminescence. In agreement with the result of flow cytometric detection, ROS production significantly increased in a time-dependent manner upon LCMV infection from SphK2-deficient BMNs both with (Fig. 4E) or without PMA stimulation (Fig. 4F). The data suggests that SphK2-deficient neutrophils are significantly activated and likely less immune suppressive during LCMV infection which is in line with reduced CD244 surface expression (Fig. 3).

**Fig. 4.**
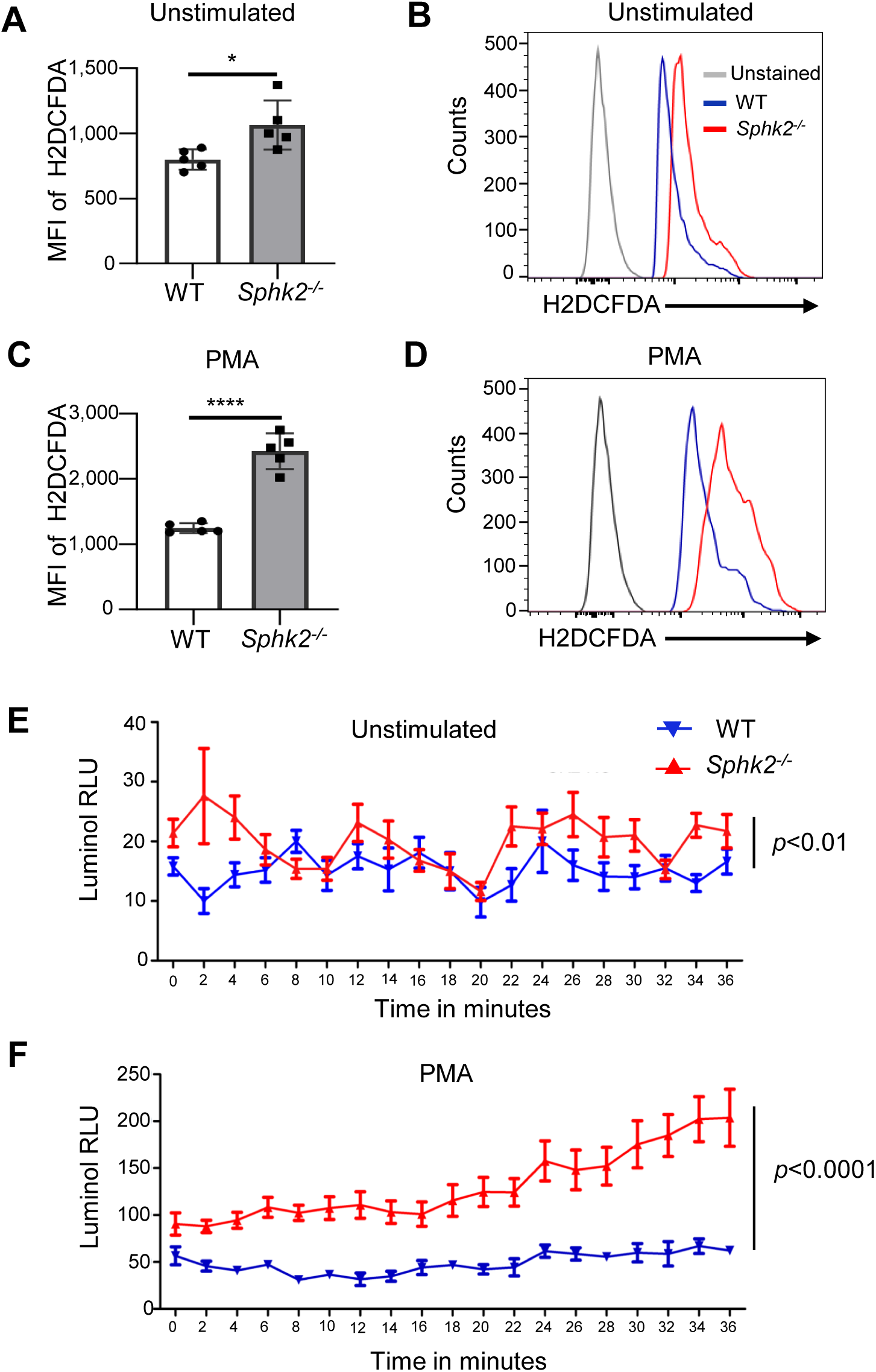
***Sphk2^-/-^* neutrophils produce significantly higher ROS upon LCMV Cl 13 infection** BMNs were isolated from LCMV Cl 13 infected WT and *Sphk2*^-/-^ mice (n = 5/group) at 3 dpi and cultured in the absence (**A-B**) or presence of 50 nM PMA for 30 min at 37^°^C CO2 incubator (**C-D**). To detect the intracellular ROS, neutrophils were incubated with H2DCFDA intracellular ROS detection probe (**A-D**) at 4^°^C for 30 min in dark. The mean fluorescence intensities (MFIs) of H2DCFDA were detected by flow cytometry (**A** and **C**) and representative histograms are shown (**B** and **D**). Production of intracellular ROS was monitored in a time-dependent manner by the luminol method in the absence (**E**) or presence of 50 nM PMA (**F**). ****p≤0.0001, *p≤0.05, unpaired t test performed for **A** and **C** and 2-way ANOVA for **E** and **F**.

### Depletion of neutrophils during chronic LCMV infection improves virus-specific T cell functions and enhances virus clearance

Neutrophils can cross-talk with various immune cells, including T cells, and orchestrate activation, proliferation, and differentiation in certain conditions (24, 34). To reveal the role of neutrophils during chronic LCMV infection, we depleted neutrophils using αLy6G treatment on day 15, 18 and 21 post-infection, followed by analysis of anti-viral T cell responses and virus clearance (Fig. 5A). Flow cytometry analysis showed that αLy6G treatment efficiently depleted neutrophils (Supplementary Fig. S6). Functionally, neutrophil depletion enhanced the anti-viral T cell response. Depletion resulted in increased IFN-γ production, as well as increased LCMV GP_66-77_ (GP66)-specific CD4^+^ T cells (GP66 tetramer^+^CD4^+^ T cells) (Fig. 5B). Similarly, we observed significantly increased LCMV GP_33-41_ (GP33)-specific CD8^+^ T cells (GP33 tetramer^+^CD8^+^ T cells) and production of intracellular IFN-γ and granzyme B (GZMB) from CD8^+^ T cells (Fig. 5C). The increase in T cell functions following treatment with anti-Ly6G antibody was associated with significantly accelerated viral clearance in serum compared to the isotype control treatment (Fig. 5D). Collectively, these results confirm that neutrophils exert an immune suppressive effect on virus-specific T cells which help promote LCMV persistence.

**Fig. 5.**
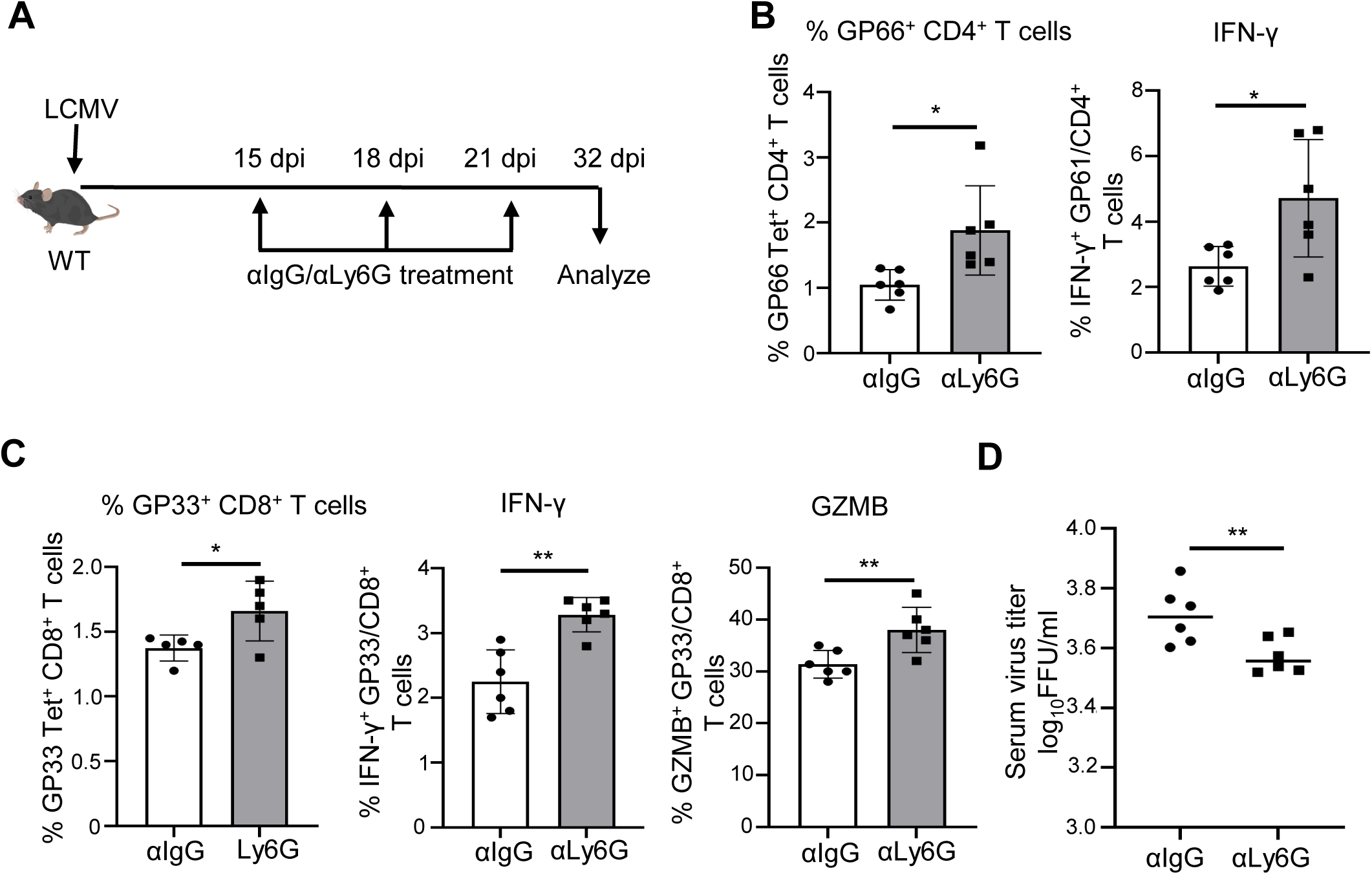
**Depletion of neutrophils during chronic virus infection restores virus-specific T cell functions and enhances virus clearance.** WT mice (n = 5-6/group) were infected with LCMV Cl 13, and neutrophils were depleted by injecting mice with 250 µg anti-Ly6G (clone 1A8) antibody via the i.p. route at 15, 18 and 21 dpi (**A**). On day 32, splenocytes were harvested. The percentage of virus-specific GP_66-77_ (GP66) tetramer^+^CD4^+^ T cells and production of intracellular IFN-γ from splenic CD4^+^ T cells upon stimulation by peptides GP_61-80_ (GP61) (**B**). The percentage of virus-specific GP_33-41_ (GP33) tetramer^+^CD8^+^ T cells in the spleen and production of intracellular IFN-γ and granzyme B (GZMB) upon stimulation by GP33 peptide (**C**) from CD8^+^ T cells was analyzed. The virus titer in the serum on 32 dpi was measured by FFU assay (**D**). **p≤0.01, *p≤0.05, bidirectional, unpaired Student’s *t*-test.

### Adoptively transferred *Sphk2*^-/-^ neutrophils promote LCMV-specific T cell response and accelerate virus clearance

The above results demonstrate the immune suppressive effect of neutrophils on virus-specific CD4^+^ and CD8^+^ T cells during chronic LCMV infection. However, SphK2-deficient neutrophils showed significantly decreased inhibitory receptors on their surface and produced significantly increased ROS when exposed to LCMV, indicating the pro-inflammatory functions of *Sphk2^-/-^* neutrophils. To test whether SphK2-deficient neutrophils exert an immune regulatory effect on T cells during LCMV infection, we employed an adoptive transfer strategy, as shown in Fig. 6A. BMNs were isolated from LCMV Cl 13-infected WT or Sphk2^-/-^ mice at 3 dpi. The comparable purity of enriched WT and *Sphk2^-/-^* BMNs was confirmed before transfer (Supplementary Fig. S7). These BMNs were transferred to LCMV Cl 13-infected WT mice at 3 and 5 dpi. On day 8 post-infection, LCMV antigen-reactive CD4^+^ and CD8^+^ T cell responses were analyzed from the recipient mice. Mice that received SphK2-deficient neutrophils showed significantly increased virus-specific CD4^+^ (Fig. 6B) and CD8^+^ T cells (Fig. 6C). Further, the transfer of SphK2-deficient neutrophils significantly enhanced the functional properties of CD4^+^ and CD8^+^ T cells, i.e., increased IFN-γ, TNF-α, and granzyme B production, compared to the T cells of mice that received WT neutrophils. T cell analyses were performed using antigenic re-stimulation with the peptides GP_61-80_ (GP61) (Fig. 6B) or GP_33-41_ (GP33) (Fig. 6C). These results demonstrate that *Sphk2*^-/-^ neutrophils have an immune stimulatory effect on virus-specific T cells during LCMV infection in vivo.

**Fig. 6.**
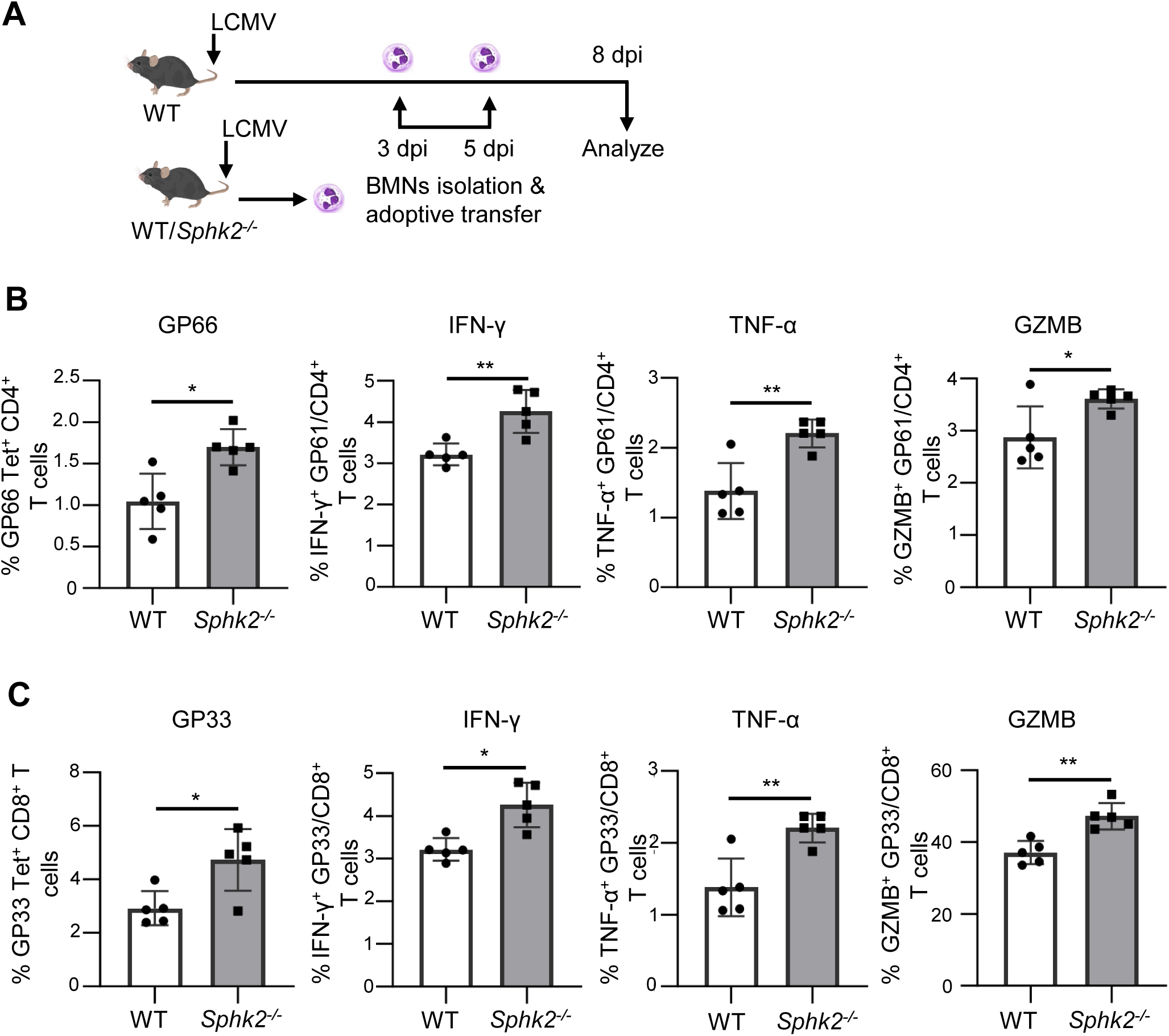
**Adoptive transfer of *Sphk2^-/-^* neutrophils significantly improves the T cell responses to LCMV infection.** BMNs were isolated from LCMV Cl 13-infected WT or *Sphk2^-/-^* mice (n = 5) at 3 dpi and then adoptively transferred to LCMV Cl 13-infected recipient WT mice (n = 5/group) at 3 and 5 dpi. The schematic presentation of the adoptive transfer strategy is depicted (**A**). On day 8 post LCMV infection, splenocytes were collected from recipient mice and CD4^+^ and CD8^+^ T cells were in vitro re-stimulated with GP61 or GP33 peptide respectively. GP66 Tet^+^CD4^+^ T cells (**B**) and GP33 Tet^+^CD8^+^ T cells (**C**) were assessed for their production of IFN-γ, TNF-α, and GZMB. **p≤0.01, *p≤0.05, bidirectional, unpaired Student’s *t*-test.

Next, we designed a similar longitudinal study to test whether these immune stimulatory effects are sustained and promote accelerated clearance of the virus. As depicted in Fig. 7A, WT or *SphK2*^-/-^ neutrophils from LCMV-infected mice were adoptively transferred at 3, 5, 12 and 19 dpi; T cell responses and virus titer were examined at 32 dpi. Similar to prior experiment (8 dpi), we observed significantly increased GP66-specific CD4^+^ and GP33-specific CD8^+^ T cells, as well as significantly increased IFN-γ and granzyme B production, at 32 dpi in SphK2-deficient neutrophil recipient mice (Fig. 7B and 7C). This is supported by significantly reduced exhaustion markers PD-1 and TIM-3 expression on the surface of both virus-specific CD4^+^ and CD8^+^ T cells (Fig. 7B and 7C). Furthermore, the virus titer in serum was significantly reduced at both 25 dpi and 32 dpi (Fig. 7D and 7E), and the virus titer in liver and kidney was decreased at 32 dpi in *Sphk2*^-/-^ neutrophil-recipient mice compared to WT neutrophil-recipient mice (Fig. 7F and 7G). These results indicate that SphK2-deficient neutrophils have immune stimulatory effects on T lymphocytes during chronic LCMV infection, which helps in overcoming T cell suppression to promote the elimination of LCMV from serum and other organs.

**Fig. 7.**
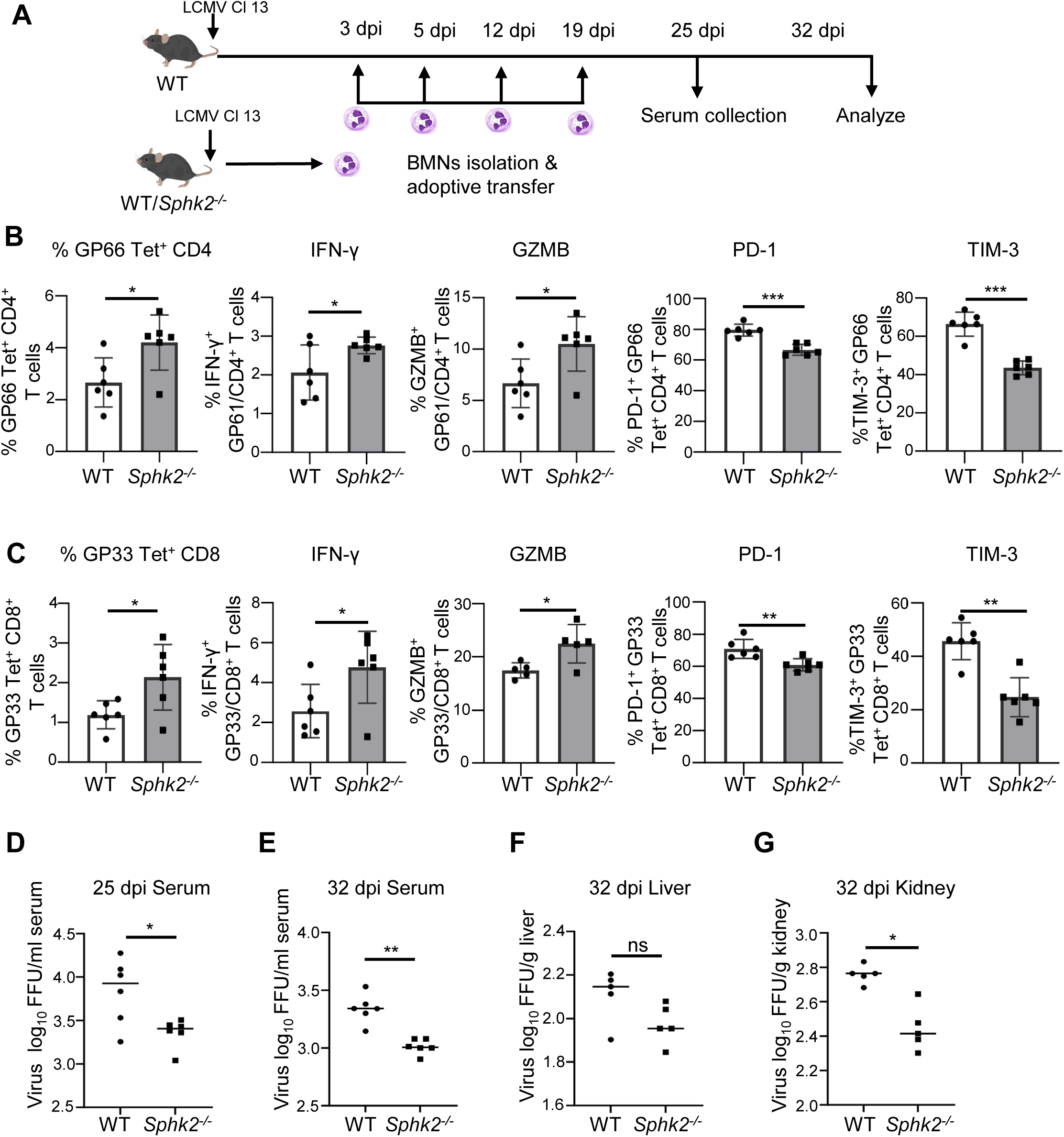
**Adoptively transferred *Sphk2^-/-^* neutrophils improve LCMV-specific T cell response and accelerated virus clearance.** BMNs from LCMV Cl 13 infected WT or *Sphk2^-/-^* mice (n = 6) were adoptively transferred to LCMV Cl 13 infected recipient mice (n=6/group) at multiple time points as shown in the workflow diagram (**A**). The number of virus-specific T cells, intracellular IFN-γ and GZMB production, expression of PD-1 and TIM-3 on GP66 Tet^+^CD4^+^ (**B**) and GP33 Tet^+^ CD8^+^ T cells (**C**) were measured by flow cytometry. The virus titers in the serum at 25 dpi (**D**) and 32 dpi (**E**), liver (32 dpi) (**F**), and kidney at 32dpi (**G**) were measured by FFU assay. ***p≤0.001, **p≤0.01, *p≤0.05, bidirectional, unpaired Student’s *t*-test.

### Innate pro-inflammatory and neutrophil activation-related genes are upregulated in *Sphk2^-/-^* neutrophils upon LCMV infection

In response to infection, a large number of neutrophils can be released into circulation. However, different environmental cues can drive the development of distinct neutrophil subsets (24, 35). The changes in the neutrophil gene expression profile during the initial phase of infection are of great interest as the accumulation of neutrophils at the site of infection and inflammation can alter the course of disease outcomes (36). Since SphK2-deficient BMNs isolated at 3 dpi displayed immune stimulatory activity, we next performed scRNA-seq of BM cells from WT and *Sphk2^-/-^* mice following 3 days of LCMV Cl 13 infection. The bone marrow cell types were annotated with cell-specific standard gene expression profile (Supplementary Fig. S8) (Fig. 8A). Based on these annotations, bone marrow neutrophils were selected for further gene expression and pathway analysis. A total of 972 transcripts were obtained from WT BMNs, of which 96 were found exclusively in WT neutrophils. Similarly, we obtained 978 transcripts from *Sphk2^-/-^* BMNs, 126 of which were restricted to *Sphk2^-/-^* and not found in WT neutrophils. Based on the expression thresholds (1.5-fold increase or decrease in expression) we obtained 116 differentially expressed genes (DEGs) between WT and *Sphk2^-/-^* neutrophils. The top 30 DEGs included genes related to antimicrobial activity (*Ngp, Ltf*, *Camp*, *Pglyrp1*, and *Ifitm3*) (37, 38), proinflammatory responses (*Pglyrp1*, *Wfdc21*, *Hmgb2*, *S100a6,*) (38, 39), cell migration and reactive oxygen species production (*S100a6*, *Cybb*, *Cmss1,* and *Hmgb2*) (40, 41), and were upregulated in *Sphk2^-/-^* neutrophils (Fig. 8B). On the other hand, genes involved in tissue repair, suppression of ROS (*Chil3, Mmp8, Malat, Trim30,* and *Trim12a*) (42–44), inhibition of neutrophil chemotaxis (*Arhgap15)* (45), antigen presentation inhibition (*Stfa2 and Stfa3*)(46) and anti-inflammatory response (*Chil3, Ifi208, and Trim30d*) (46, 47) were decreased on *Sphk2^-/-^* neutrophils (Fig. 8B). These findings emphasize that the deletion of *Sphk2* alters the gene expression levels in bone marrow neutrophils, and hence neutrophil heterogeneity upon Cl 13 infection.

**Fig. 8.**
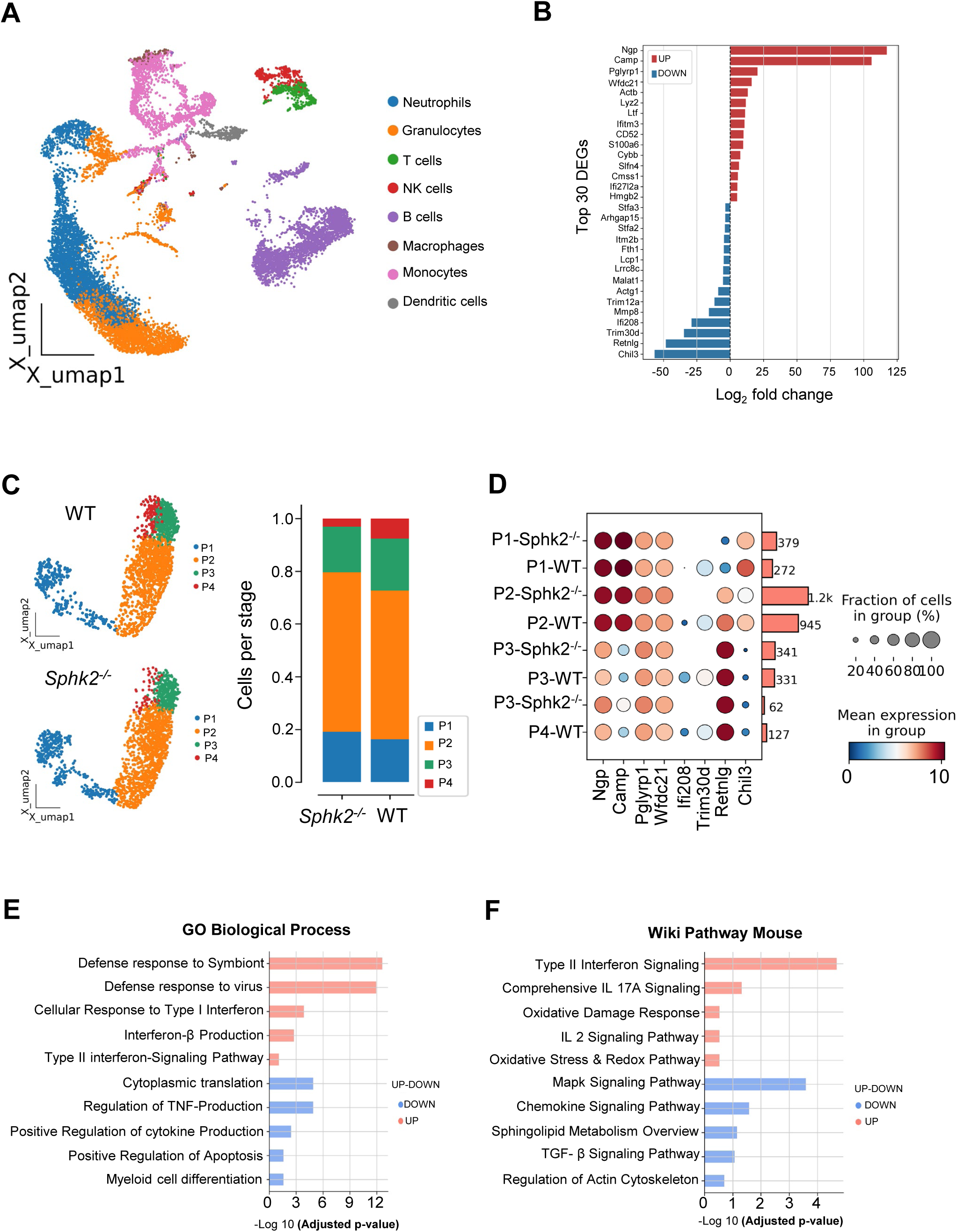
***Sphk2^-/-^* neutrophils from LCMV Cl 13 infected mice display a proinflammatory phenotype:** WT and *Sphk2*^-/-^ mice (n = 3/group) were infected by LCMV Cl 13, uninfected C57BL/6 WT mice (n = 3) were used as control. At 3 dpi bone marrows (BM) were collected, and BM cells were isolated and resuspended in PBS+0.04% BSA at a density 1 × 10^6^ cells/ml. The cell suspensions were immediately sent to the DNA core facility for single-cell RNA-sequencing. An overview of UMAP of the cell cluster derived from BM of uninfected WT or infected WT and infected Sphk2^-/-^ mice (n = 3), different color indicates distinct cell types (**A**). The log fold changes for the top 30 differentially expressed genes (DEGs) that are upregulated or downregulated in *Sphk2^-/-^* mice (**B**). UMAP of bone marrow neutrophil subclusters P1-P4 distribution and a bar graph showing the proportion of each neutrophil subcluster from *Sphk2^-/-^* and C57BL/6 WT mice (**C**). Distribution of selected topmost DEGs on neutrophil subsets of WT and *Sphk2^-/-^* mice is shown, and cell events corresponding to different subsets are depicted by the bars on the right (**D**). GO analysis of biological processes (BP) and Wiki pathways of up- and downregulated pathways in infected *Sphk2^-/-^* neutrophils compared to infected WT neutrophils (**E and F**).

To assess the distribution of the top DEGs on different neutrophil subsets, we classified neutrophils into P1-P4 sub-types based on their gene expression profile as described by Grieshaber-Bouyer, *et al* 2021 (48) (Fig. 8C) (Supplementary Fig. S9A). These 4 subtypes represent highly proliferating P1 pre-neutrophils, immature P2-P3 neutrophils, and mature P4 neutrophils. Upon infection, matured neutrophils (P4) rapidly moved into circulation, hence their proportion is decreased in bone marrow. As shown in Fig. 8C, the presence of a higher number of proliferating P1 neutrophils and decreased amounts of terminally differentiated mature P4 neutrophils were observed in *Sphk2^-/-^* mice (Fig. 8C). Taken together, this suggests elevated proliferation of neutrophil progenitors and increased egress of matured neutrophils into circulation, with eventual egress into peripheral organs (Fig. 8D). Next, we mapped top DEGs for each subtype cluster to access the distribution of the top DEGs within the subtypes (Supplementary Fig. S9B). As shown in Fig. 8D, genes involved in proinflammation were preferentially distributed in P1-P2 subtypes, while anti-inflammatory and tissue repair-related genes are distributed in P3-P4 subtypes suggesting that the phenotypic changes in neutrophils could occur as early as at 3 dpi (Supplementary Fig. S8B). Further, pathway analysis revealed changes in neutrophil functional status of *Sphk2^-/-^* mice including activation of innate immune response to viral infection, interferon signaling, oxidative damage response, decreased sphingolipid metabolism, TGF-β signaling, and apoptosis (Fig. 8E and F). In summary, these results reinforce that *Sphk2-*deficient neutrophils exhibit a pro-inflammatory phenotype during chronic LCMV infection and further support our in vivo experimental findings.

## Discussion

During chronic infection, suppressive neutrophils can develop and contribute to T cell dysfunction and virus persistence. Our study identified SphK2 as a host immune regulatory molecule crucial for neutrophil suppression in the context of chronic LCMV infection. A key role of SphK2 in the regulation of neutrophil gene expression was uncovered which supports the functional changes of neutrophils upon infection.

The depletion of Ly6G^+^ cells in WT mice during LCMV Cl 13 infection increased virus-reactive T cell immunity, contributing to lowered viral burden (Fig. 5). These results indicate that Ly6G^+^ neutrophils acquire the immune suppressive functions upon infection which contributes to T cell dysfunction and LCMV persistence. Previous studies were conducted to deplete neutrophils using anti-Gr-1 antibody (RB6-8C5) during chronic LCMV Cl 13 infection (25). However, this antibody was reported to deplete other immune cells, including monocytes and eosinophils, in addition to neutrophils (49). Therefore, it was unclear whether neutrophils were indeed responsible for the regulation of T cell immunity caused by Gr1^+^ cell depletion in the prior study. As such, our study utilized anti-Ly6G antibody (1A8) treatment, which is known to specifically deplete neutrophils, to further demonstrate that neutrophils retain the immune suppressive function to impact T cell exhaustion and chronic LCMV infection. During acute infection, neutrophil numbers typically peak within the first 24 hours and gradually subside. However, LCMV Cl 13 spreads systemically and establishes viral persistence, accompanied by continuous neutrophil responses. Interestingly, SphK2-deficient mice showed significantly increased neutrophils in multiple organs upon LCMV infection which was sustained when assessed at 14 dpi (Supplementary Fig. S1). Upon LCMV Cl 13-infection, SphK2-deficient neutrophils displayed more pro-inflammatory phenotypes, as evidenced by the increased ROS production and decreased CD244 expression. The increased inflammatory neutrophilic response was shown to contribute to the viral immune pathogenesis of *Sphk2^-/-^*mice as neutrophil depletion partially improved the viability of LCMV-infected *Sphk2^-/-^* mice. The reasons for the partial rescue of LCMV Cl 13 infected *Sphk2^-/-^* mice by neutrophil depletion are currently unknown. Neutrophils are a heterogeneous population; not all are immune suppressors (50). It is possible that the antibody-mediated depletion transiently removed all the neutrophils, but that some neutrophils may benefit virus clearance or tissue repair, thereby compromising the protective effects of the depletion (51). Further, immune cells other than neutrophils might also promote viral immune pathogenesis (23) and SphK2 may display immune regulatory functions in multiple immune cell types.

Since both suppressive and immune stimulatory neutrophils express many common phenotypic markers on their surfaces, it is difficult to differentiate them by flow cytometry. However, a few studies have used SLAMF receptor CD244 to differentiate suppressive neutrophils from pro-inflammatory neutrophils (30, 31); targeted deletion of CD244 on myeloid cells improved CD8^+^ T cell effector functions (52, 53). We found significantly reduced expression of CD244 on SphK2-deficient neutrophils during the early stage of infection (Fig. 3) (Supplementary Fig.S4). Further, the adoptive transfer experiment proves the neutrophil-intrinsic regulation of CD244 by SphK2 (Fig. 3F). CD244, also known as 2B4 or SLAMF4, was also revealed as a marker for T cell exhaustion, as its expression significantly increased on exhausted T cells during LCMV 13 infection(15). CD244 blockade along with anti-PDL-1 antibody treatment synergistically enhanced the antiviral response of CD8^+^ T cells upon LCMV Cl 13 infection, while CD244 blockade alone produced a moderate increase in the CD8^+^ T cell functions (54). It is unclear if the blockade of CD244 affected the function of suppressive neutrophils in the prior studies. Blocking of CD244 on pan-immune cells may not be a prudent approach because CD244 could act as either an inhibitory or stimulatory receptor depending on the cell types, receptor density, and availability of downstream intracellular signaling molecules (53). However, CD244 on neutrophils transmits inhibitory signaling, suggesting that neutrophil-specific targeting of CD244 could be a better strategy.

Our scRNA-seq reveals differences in the neutrophil gene expression profile at the subpopulation level between WT and *Sphk2^-/-^* mice. Genes involved in opposite pathways (tissue repair vs tissue damage) diverge at the early developmental stages. These subsets egress from bone marrow at their current stage, after which they may develop into corresponding subtypes and perform their functions accordingly (55). *Chil3, Mmp8, Ifi208, Trim30d* and *Trim12a,* which are involved in tissue remolding and repair and are therapeutically targeted against inflammatory diseases in an experimental setting (56), are significantly increased in WT neutrophils (Fig. 8B). Most importantly, *Ifi208*, *Trim30d* and *Trim12a* are undetectable in *Sphk2^-/-^* neutrophils (Fig. 8D and Supplementary Fig. S9B). *Ifi208* (*Pydc3*) belongs to the interferon inducible family p200 and has been shown to inhibit the inflammasome function (47); *Trim30* was reported to inhibit cell proliferation in lymphoid cells (57), whereas in myeloid cells it downregulates ROS production, NLRP3 inflammasome activation, and decreases the influx of neutrophils (58). *Trim12a* shares high sequence homology with *Trim30*, but the functional studies of this gene are very limited (59). Although a limited number of studies have demonstrated the immune regulatory functions of these genes, detailed studies on their regulation of neutrophil functions still need to be explored. A better understanding of SphK2’s regulation of neutrophil function will be helpful for designing advanced immune therapeutics against chronic infection as well as inflammatory conditions, which necessitate further studies on the role of SphK2 and its downstream pathway during persistent virus infection associated with immune suppression or immunopathologic inflammation.

## Materials and methods

### Sex as a biological variable

We included both male and female C57BL/6 (WT) and C57BL/6 *Sphk2*^−/−^ mice in this study, and no difference between sex were observed and the results were pooled.

### Mice

All mice used in this study were male and female C57BL/6 (WT) (the Jackson Laboratory) and C57BL/6 *Sphk2*^−/−^ mice (provided by Richard Proia, NIH, Bethesda, Maryland, USA) aged between 6 to 10 weeks at the beginning of the study. Mice were either bred and maintained in a closed breeding facility according to institutional guidelines or purchased from Jackson Laboratory. The animal studies were approved by the Animal Care and Use Committee of the University of Missouri-Columbia.

### Virus titration and infections

Lymphocytic choriomeningitis virus (LCMV) strains, LCMV Armstrong (Arm) and clone 13 (Cl 13) were propagated on BHK cells (60). LCMV titers were determined by plaque-forming unit (PFU) or focus-forming unit (FFU) assay on Vero E6 cells as described elsewhere (23, 61). Mice were infected with 2 × 10^6^ FFU of LCMV Cl 13 via the intravenous (i.v.) route (62). For LCMV Arm experiments, mice were infected with 2 × 10^5^ FFU of LCMV Arm via the intraperitoneal (i.p.) route. 6-8 weeks old uninfected WT and *Sphk2^−/−^* mice were used as background controls in all in vivo experiments.

### Enrichment of bone marrow neutrophils (BMNs)

Hind legs were collected in ice-cold HBSS with 2% FCS and placed on ice. All steps were carried out aseptically in the laminar air chamber, and samples were placed on ice. Samples briefly dipped in 70% ethanol for 20-30 seconds. Femurs and tibias were separated, and bone marrow was flushed with 1 ml of ice-cold HBSS+2% FBS using 26-gauage syringe. The debris were removed by passing the cell suspension through a 40 µM cell strainer. Cells were collected in 50 ml falcon tube and centrifuged at 300 x g at 4^°^C for 5 min. RBCs were lysed by resuspending the pellet in 10 ml 0.2% NaCl for 20-25 seconds and neutralized by addition of equal volume (10 ml) 1.6 % NaCl solution. Tubes were centrifuged as above, and pellet was resuspended in HBSS + 2% FBS buffer. To separate neutrophils from other mononuclear cells, 2 ml Histopaque 1119 (Sigma, St.Louis, MO) was taken in 15 ml falcon tube and on the top of this layer 3 ml Histopaque 1083 (Sigma, St.Louis, MO) was added slowly. On the top of this Histopaque gradient, 3 ml bone marrow cell suspension was gently overlaid and centrifugation at 872 × g for 25 min at room temperature (RT). The upper mononuclear cells band was discarded and the neutrophil rich band present at the junction between Histopaque 1119 and 1083 was collected in 10 ml RPMI medium and washed twice as mentioned above. The purity of the neutrophils was confirmed by neutrophil specific cell surface markers staining using anti-mouse CD45, CD11b, and Ly6G antibodies by flow cytometry acquisition and analysis and used for downstream processing.

### Isolation of lymphocytes

Splenocyte suspensions were obtained by mashing the spleen through a 40 μm cell strainer. Cells were collected and pelleted by centrifuging at 300 × g for 5 min at RT. RBCs were lysed using 1 × RBC lysis buffer (Sigma), and cells were washed twice as mentioned above and finally resuspended in T cell medium (RPMI containing 10 % FBS, 1% antibiotics, 1% non-essential amino acids, 1 mM HEPES and 1 mM sodium pyruvate).

### Isolation of kidney cells

Kidney capsules were removed and chopped into small pieces and transferred to a 5 ml serum-free RPMI medium containing 0.2 mg/ml collagenase D (Gibco) and 0.1 mg/ml DNase I (Sigma). The content was transferred to gentleMACS dissociation C tube (Miltenyi Biotech) and homogenized per manufacture’s protocol. The samples were incubated in 37^°^C water bath for 30 min. The samples were again homogenized and passed 100 µm cell strainer and centrifuged at 300 × g for 5 min at RT. The pellet was resuspended in 1 ml RBC lysis buffer and incubated for 3 min at RT, and RPMI medium was added to stop the lysis and centrifuged as above. Cells were resuspended in HBSS containing 2% FBS and overlaid on 33 % Percoll and centrifuged at 300 x g for 15 min at RT. The upper debris was removed, and pellets were resuspended in RPMI medium and used for downstream processing.

### Neutrophil depletion

Neutrophils were depleted by administering 250 µg anti-Ly6G antibody (Clone 1A8, Leinco Technologies, St. Louis, MO) via the i.p. route. Control groups were treated similarly with IgG isotype control antibody, IgG2a (Leinco Technologies St. Louis, MO), on indicated days as described in the respective figure legends.

### Adoptive transfer of neutrophils

For neutrophil adoptive transfer experiments, WT and *Sphk2*^−/−^ mice were infected with LCMV Cl 13, and BMNs were isolated at 3 dpi as described above. Purity (80-90%) of the neutrophil was assessed by flow cytometry prior to the adoptive transfer of 2 x 10^6^ neutrophils into WT mice by i.v. injection on the indicated days.

### Adoptive transfer of CFSE-stained neutrophils

BMNs were isolated from uninfected WT and Sphk*2*^−/−^ mice and were incubated with 5µg/ml CFSE (ThermoFisher) at 37^°^C in the dark for 15 min. Cells were then washed twice with RPMI medium and finally resuspended in 1x DPBS. Subsequently, these CFSE-stained neutrophils (2 ×10^6^) were adoptively transferred to LCMV Cl 13-infected wild-type mice at 3 dpi. After two days of transfer, spleen and kidney samples were collected to assess the expression of CD244 on adoptively transferred donor neutrophils in recipient mice.

### Reactive Oxygen Species (ROS) quantification

BMNs were isolated from LCMV Cl 13-infected WT and *Sphk2*^−/−^ at 3 dpi. 1 × 10^5^ cells/well were seeded in a 96-well plate and cultured in CO_2_ incubator for 30 min at 37°C in the presence or absence of 50 nM PMA. Cells were washed and resuspended in 100 µl FACS staining buffer containing 5 µM H2DCFDA. The plate was incubated at 4°C for 30 min in dark. Cells were washed twice with FACS stain buffer and samples were acquired using LSR Fortessa X420 flow cytometry. The mean fluorescence intensity (MFI) was analyzed by FlowJo software. ROS production in a time-dependent manner was measured using a luminol-enhanced chemiluminescence assay. Briefly, 1 ×10^5^ cells/well were seeded in 96 well plate and cultured in CO_2_ incubator for 15 min at 37^°^C in the presence 5µM luminol. After incubation, 50nM PMA was added to the designated wells and intracellular chemiluminescence was read from the top of each well at 2 min intervals for 40 min by microplate reader (CLARIOstar^®^ *Plus, BMG LABTECH*).

### Flow cytometric analysis

The surface antibody staining was carried out on a 96-well V-bottom plate. 1 x 10^6^ cells/well were resuspended in 50 µl FACS staining buffer containing appropriate antibody concentration. For flowcytometry analysis, singlet cells were selected and CD45 positive leukocytes were used for further analysis. LCMV GP_33-44_–specific CD8^+^ T cells were identified using fluorochrome-linked GP_33-41_ tetramers, and LCMV GP_66-77_-specific CD4^+^ T cells were identified using fluorochrome-linked GP66 tetramers, which were provided by the NIH Tetramer Core Facility (Emory University, Atlanta, Georgia, USA). For intracellular cytokine staining, lymphocytes were cultured in the presence of 10 μg/mL of brefeldin A (Sigma, St.Louis, MO, USA) and 2 μg/mL GP33 (KAVYNFATC), 5 μg/mL GP_61-80_ (GP61, GLNGPDIYKGVYQFKSVEFD) peptide for 6 hours, followed by fixation, permeabilization, and staining with the indicated antibodies. Samples were run on LSR Fortessa X-20 (BD Biosciences) or Cytek Aurora spectral analyzer (Cytek Biosciences) and analyzed with FlowJo software. Antibodies used in this study can be found in the appendix section.

### Determination of virus titers

Tissues from the liver, kidney, and spleen along with serum were harvested from infected mice at the time indicated. Tissues were homogenized using a Bead Beater with 1.0 mm diameter Zirconia/Silica beads (BioSpec Products). LCMV titers were determined by FFU or PFU assay on Vero cells as performed earlier(23).

### Single-cell RNA-sequencing and bioinformatic analysis

Bone marrows were collected from uninfected WT and LCMV Cl 13-infected WT and *Sphk2*^−/−^ mice at 3 dpi. Bone marrow cells were isolated as mentioned above with little modification in the isolation and suspension buffer. Cells were passed through a 40 µM cell strainer twice to remove debris and aggregates. Total bone marrow cell suspensions were resuspended in PBS+0.04% BSA buffer. Finally, 1×10^6^ cells/ml with >85% viable bone marrow cell suspensions were prepared and immediately handed over to the university DNA core facility for scRNA-seq.

### 10x Genomics Single Cell 3’ RNA-Seq Library Preparation Method

Libraries were constructed by following the manufacturer’s protocol with reagents supplied in 10× Genomics Chromium Next GEM Single Cell 3′ Kit v3.1. Briefly, cell suspension concentration and viability were measured with a Cellometer K2 (Revvity) stained with an acridine orange/propidium iodine dye mix (Invitrogen). Subsequently, cell suspension (targeting 10,000 cells) combined with reverse transcription master mix was loaded on a Chromium Next GEM chip G along with gel beads and partitioning oil to generate gel emulsions (GEMs). GEMs were then transferred to a PCR strip tube, followed by the reverse transcription, which was performed on a Veriti thermal cycler (Applied Biosystems) at 53°C for 45 minutes. cDNA was amplified for 11 cycles and purified using AxyPrep MagPCR Clean-up beads (Axygen). cDNA fragmentation, end-repair, A-tailing and ligation of sequencing adaptors were performed according to the manufacture’s specifications. Library concentration was measured with a Qubit HS DNA kit (Invitrogen) and fragment size was measured on a 5200 Fragment Analyzer (Agilent). Libraries were pooled and sequenced on a NovaSeq X (Illumina) to generate 50,000 reads per cell with a sequencing configuration of 28 base pair (bp) on read 1 and 100 bp on read 2.

### scRNA-seq data processing

Single-cell RNA sequencing (scRNA-seq) data from mouse bone marrow were processed using the CellRanger toolkit (v8.0.1) with alignment to the GRCm39 mouse reference genome. Only reads that were uniquely aligned, non-PCR duplicates, and associated with valid cell barcodes and unique molecular identifiers (UMIs) were used to construct the initial 3′ gene-by-cell matrix, comprising 30,500 cells.

To correct for ambient RNA contamination and barcode swapping, we applied CellBender (v0.3.2) (63). Cells were retained if they passed a false discovery rate (FDR) threshold of <0.01, adjusted using the Benjamini–Hochberg method. Additional quality filters were imposed to exclude low-quality cells: (1) total UMI count >1,000, (2) number of detected genes >500, and (3) proportion of mitochondrial transcripts <10%.

Potential doublets were further filtered using a cluster-level approach. DoubletDetection (v4.3) was used to compute a doublet score for each cell (64). Cells were grouped into clusters per sample by computing the top 50 principal components from the 3,000 most variable genes, followed by construction of a k-nearest neighbor graph via the pp.neighbors function, and clustering with the Louvain algorithm implemented in Scanpy (v1.11.0) (65, 66). Median-centered, MAD-scaled doublet scores were calculated for each cluster to identify and remove likely doublets.

After all quality control procedures, 19,004 high-confidence cells from three biological samples (nine mice) were retained for downstream analyses. For each retained cell, both raw UMI counts and log₂-transformed normalized expression values were computed.

### Integrated analysis of single-cell datasets

To account for inter-sample variability and batch-associated confounders, we utilized the scvi-tools framework to perform integrated analysis of single-cell transcriptomes (67). This approach applies single-cell variational inference (scVI), a deep generative modeling technique built upon variational autoencoders (VAEs), to learn a unified latent representation of gene expression profiles while controlling batch effects.

In the scVI model, each cell is encoded into a low-dimensional latent variable that is conditioned on known batch annotations. The generative decoder then reconstructs the observed expression levels using this latent embedding in combination with the batch covariate. By explicitly incorporating batch as a covariate during training, scVI disentangles biological variation from technical artifacts and captures the intrinsic structure of the data across samples.

During model inference, batch labels can be marginalized out to yield batch-harmonized latent embeddings, which support integrated downstream analyses such as dimensionality reduction, clustering, and trajectory inference. These corrected representations offer a probabilistically grounded and scalable solution for batch correction, particularly well-suited for large and heterogeneous single-cell datasets.

### Visualization and signature-based cell classification

Major immune cell types were identified using the Scanpy Python package. We first selected 3,000 highly variable genes (HVGs), followed by principal component analysis (PCA) to compute the top 50 components for dimensionality reduction. Prior to downstream analysis, the percentage of mitochondrial transcripts was regressed out, and all gene expression values were scaled to unit variance. A k-nearest neighbor graph was constructed using the pp.neighbors function, and clustering was performed with the Leiden algorithm at a resolution of 1, resulting in 26 transcriptionally distinct clusters. The structure of the cellular landscape was visualized using UMAP for two-dimensional embedding.

Initial annotation of major immune cell types was performed using SingleR (v2.10.0) (68), referencing the MouseRNAseqData database from Celldex (v1.18.0). To further refine these annotations, we computed gene signature scores using the tl.score_genes function in Scanpy, based on canonical marker genes. Neutrophils were characterized by high expressions of *Ly6g, Itgam, Cxcr2, Csf3r, S100a8, Il1r2, Trem1, Ceacam1, Hp,* and *Hdc*. Neutrophils were defined as cells exhibiting both *Ly6g* expressions greater than zero and a positive neutrophil gene module score. Macrophages expressed *Mrc1, Cd68*, and *Adgre1*, while monocytes were marked by *Plac8, Psap,* and *Ccr2*. Dendritic cells were identified by expression of *Km0* and *Flt3*. T cells were annotated based on *Cd3d, Cd3e*, and *Trbc2*, whereas B cells expressed *Cd79a*, *Cd79b*, and *Pxk*. Natural killer cells showed high levels of *Klrd1, Klrk1,* and *Nkg7*, and granulocytes were distinguished by *Csf3r*, *Clec4d,* and *Cxcr2* expression.

To investigate transcriptional heterogeneity within the neutrophil population, we conducted a second round of clustering exclusively on neutrophil-annotated cells. Gene module scores were calculated for subpopulation classification based on curated markers as described by Grieshaber-Bouyer, et al 2021(48). One subpopulation, designated as p1, was defined by elevated expression of *Cebpe, Hmgb2, Chil3, Ngp, Arhgdib, Calm2, mt-Co1, Lcn2, Lyz2, Wfdc21, Cybb, Ly6c2, Ly6g, Cd177, Serpinb1a, Pglyrp1,* and *Prdx5*. A second group, p2, was enriched for *Ltf, Lcn2, Lyz2, Wfdc21, Ifitm6, Anxa1, Mmp8, Cybb, Dstn, Ly6c2, Ly6g, Cd177, Prdx5, Lgals3, Mmp9, Pglyrp1,* and *Mcemp1*. The third subpopulation, p3, showed elevated levels of *Mmp8, Lgals3, Mcemp1, Retnlg, S100a6, Prr13, Fth1, Ccl6, Msrb1,* and *H2-D1*. Finally, the p4 subset was characterized by expression of *Wfdc17, Ifitm1, Ifitm2, Btg1, Fxyd5, Srgn, Malat1, Dusp1, Rps27, Jund, Fth1, Msrb1, Csf3r, Junb,* and *H2-D1*.

Altogether, we identified eight major immune cell types and four transcriptionally distinct neutrophil subpopulations. Differentially expressed genes across clusters and neutrophil subsets were identified using the tl.rank_genes_groups function in Scanpy, applying the Wilcoxon rank-sum test and controlling the false discovery rate using the Benjamini–Hochberg procedure. Genes with an adjusted P-value below 0.05 were considered statistically significant.

The complete and detailed scRNAseq data can be found in the GEO database (GSE303763).

### Pathway enrichment analysis

To investigate the functional relevance of genes identified as cluster-specific markers, we performed gene set enrichment analysis using GSEApy (v1.1.8) (69). Differential expression results from the *Sphk2(+)* neutrophil population under WT treatment were used to define gene sets of significantly upregulated and downregulated transcripts. These gene sets were queried against multiple curated databases, including Enrichr, the Gene Ontology (GO) Biological Process, and WikiPathways signaling collections (70). This analysis enabled the identification of biological processes and signaling pathways significantly associated with the transcriptional response of neutrophils to treatment, providing insight into the molecular programs regulated by Sphk2 activity.

### Statistical analysis

All error bars represent the mean ± SEM, and averages were compared using a bidirectional (2-tailed), unpaired Student’s t-test unless otherwise indicated. In the case of different sample sizes, an unequal variances t-test was employed. For virus titers, an unpaired two-tailed t-test was used to account for nonparametric viral clearance. Graph constructions and statistical analyses were performed using Prism 10 (GraphPad). For time-dependent ROS production between wild-type and *Sphk2^-/-^* mice, a 2-way ANNOVA test was performed. For the mouse survival study, the Kaplan-Meier test was used to determine statistical significance between different treatments. A p-value less than or equal to 0.05 was considered statistically significant for this analysis. For scRNA-seq data, a q-value (FDR) of less than 0.05 was used, while for GSEA pathway analyses, a q-value (FDR) of less than 0.1 was considered significant.

## Acknowledgements

The study was supported by NIH/NIAID R01AI153076 and R01AI162631 (B.H.) and University of Missouri School of Medicine funds (LHA). We thank Dr. Richard Proia (NIH/NIDDK), Kelley Argraves and Lina Obeid (MUSC) for providing *Sphk2^-/-^* mice. We sincerely thank Regina Wamsley for her technical assistance during this study. We thank Dr. Michael Oldstone (Scripps) for LCMV ARM & LCMV Cl 13 virus strains. We also thank the NIH Tetramer Core Facility for the kind provision of GP33 and GP66 tetramers.

