## Supplementary figures and table for "Sphingosine kinase 2 suppresses neutrophil responses to promote viral persistence while attenuating immune pathology"

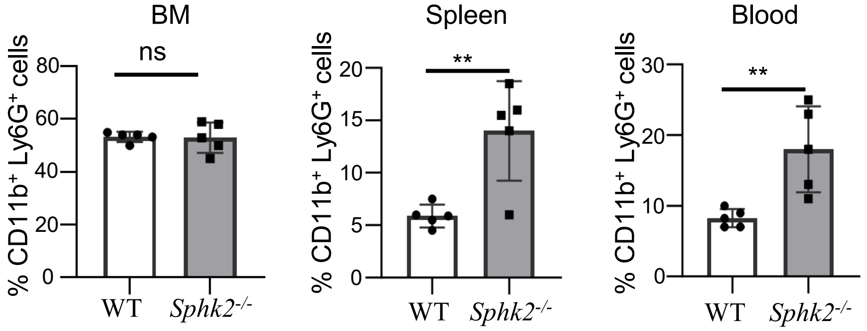


**Fig. S1. LCMV Cl 13 infection induces enhanced and sustained neutrophils in *Sphk2$^{-/-}$* mice.** WT and *Sphk2$^{-/-}$* mice (n = 5/group) were infected with LCMV Cl 13. Mice were euthanized at 14 dpi, and organs were collected. The percentage of CD11b$^{+}$Ly6G$^{+}$ neutrophils in bone marrow (BM), spleen, and blood of LCMV Cl 13-infected WT and *Sphk2$^{-/-}$* mice was quantified using flow cytometry. **p≤0.01, *n.s.* not significant, bidirectional, unpaired Student's *t*-test.

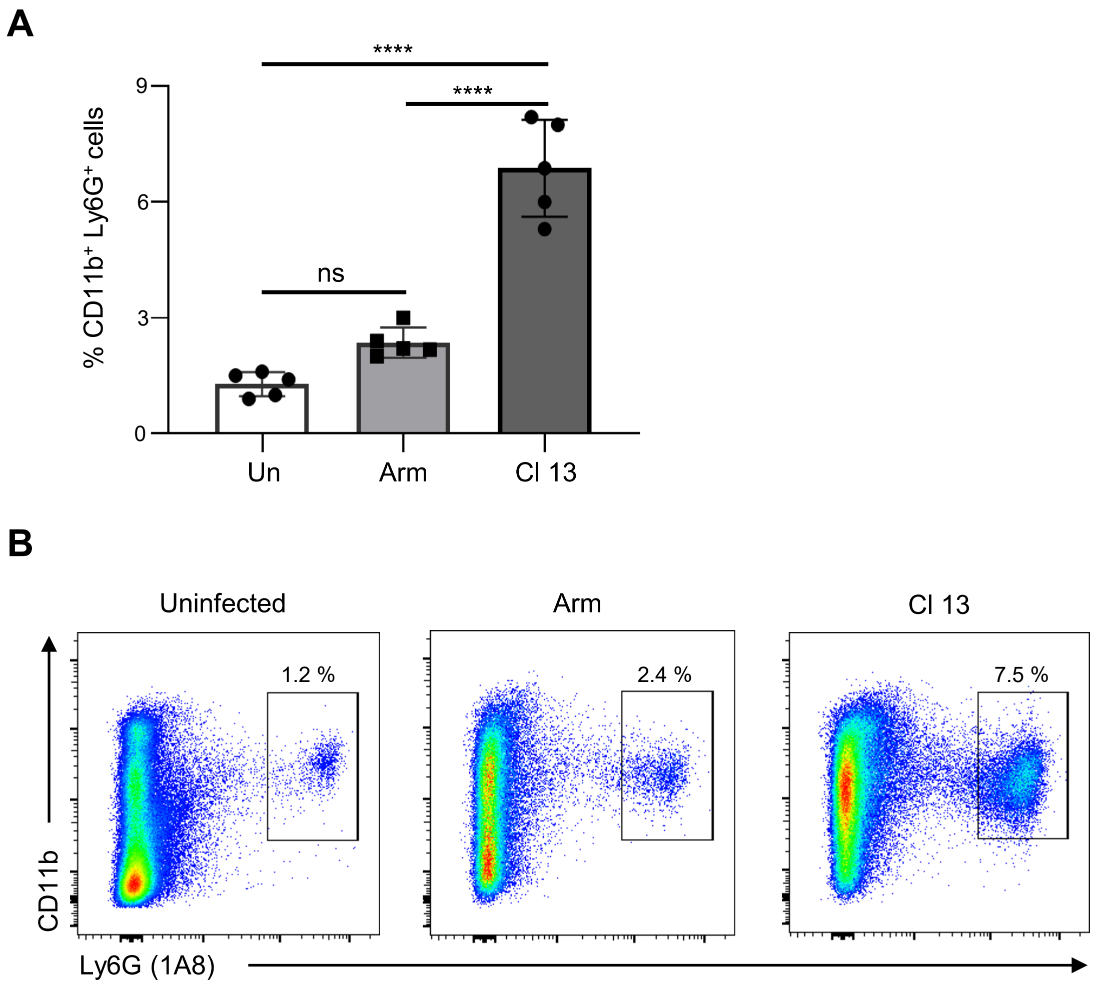


**Fig. S2. LCMV Cl 13 induces higher neutrophils compared to LCMV Arm.** WT (n = 5/group) were infected with LCMV Arm or Cl 13. At 10 dpi, mice were euthanized and splenocytes were isolated. The percentage of $CD11b^+Ly6G^+$ neutrophils from uninfected control, LCMV Arm and Cl 13 infected group was quantified by flow cytometry (**A**). The representative flow cytometric dot plot of one sample from each group is depicted (**B**). ****$p \leq 0.0001$, ns, not significant. 1-way ANOVA was performed to find the significant difference between the groups.

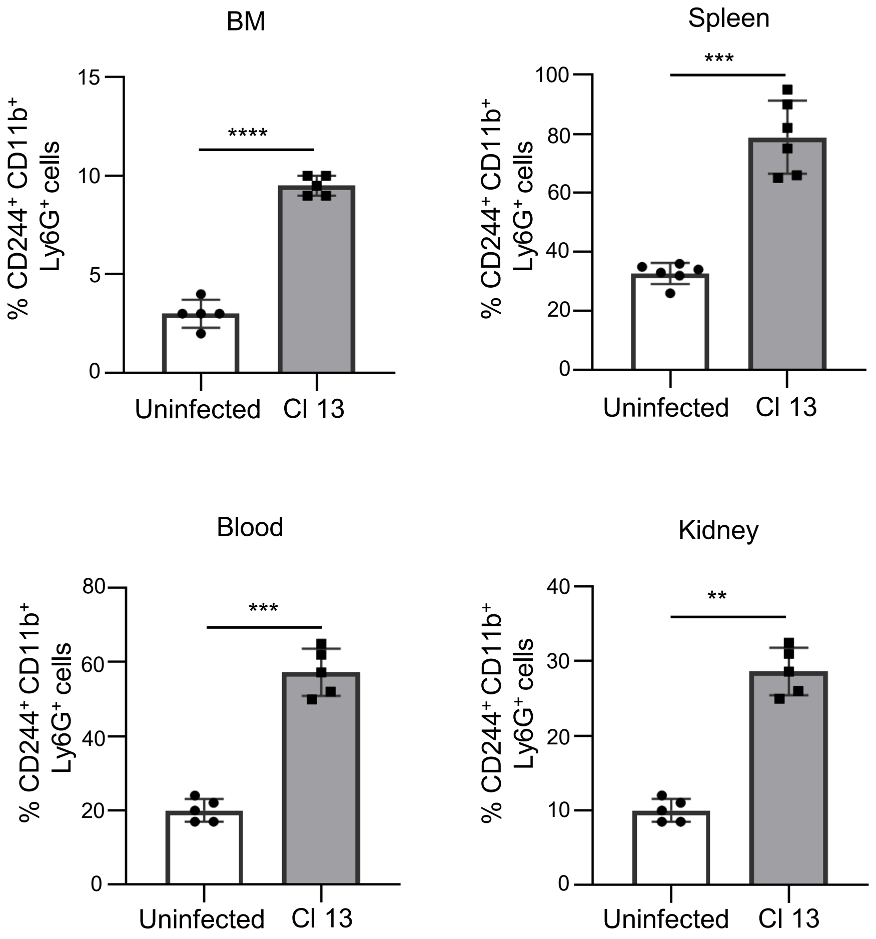


**Fig. S3. LCMV Cl 13 infection induces significantly increased $CD244^+$ neutrophils in different organs.** WT mice (n = 5-6) were uninfected (control) or infected with LCMV Cl 13, and at 3 dpi mice were euthanized and organs were collected. The percentage of $CD244^+$ neutrophils in different organs was quantified by flow cytometry, and the percentage levels were compared with those of uninfected C57BL/6 WT mice. ****$p \leq 0.0001$, ***$p \leq 0.001$, **$p \leq 0.01$, bidirectional, unpaired Student's *t*-test.

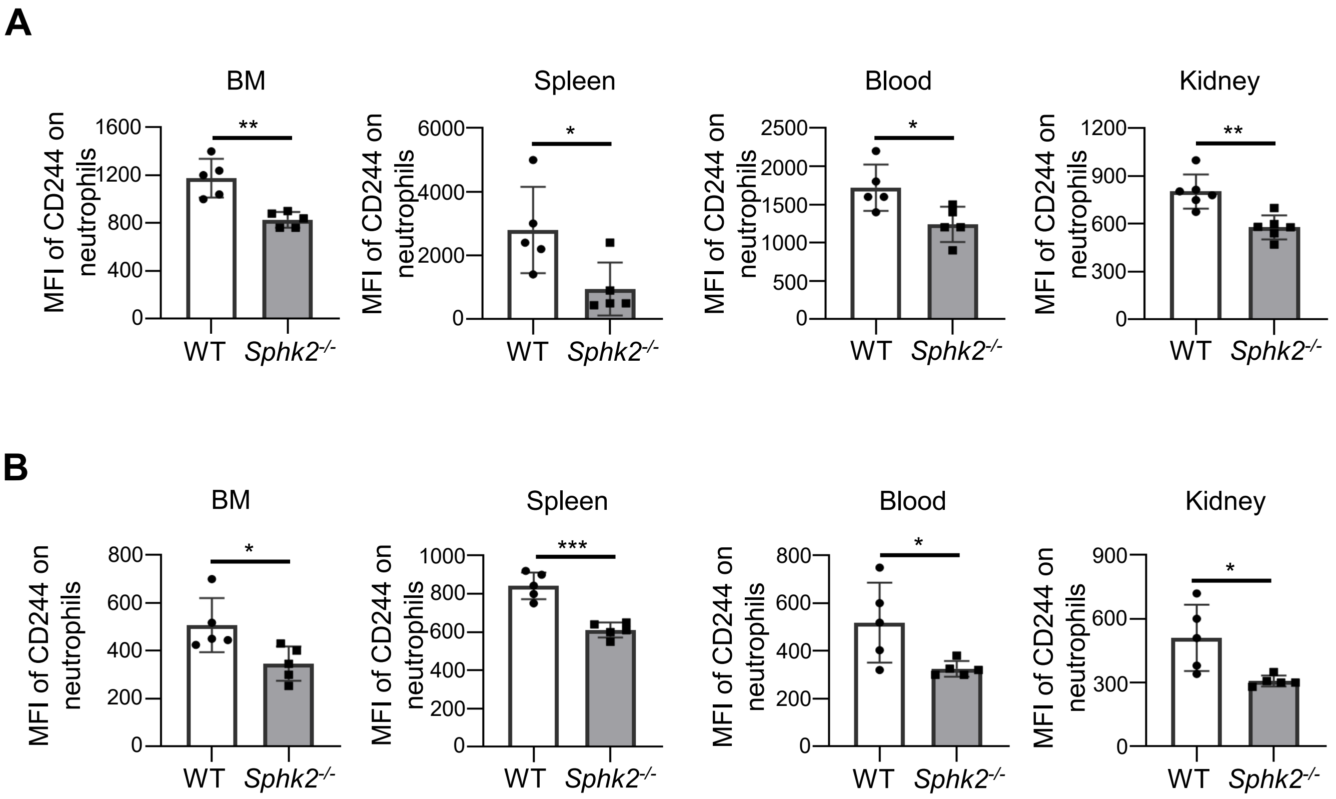


**Fig. S4. SphK2 deficiency reduces the expression levels of CD244 on neutrophils during LCMV Cl 13 infection.** WT and *Sphk2*$^{-/-}$ mice (n = 5/group) were infected with LCMV Cl 13, and at 3 and 8 dpi, mice were euthanized, and organs were collected. The geometric mean fluorescence intensity (MFI) of CD244 on neutrophils in multiple organs at 3 dpi (A) and 8 dpi (B) were measured by flow cytometry, and the expression level of CD244 were compared between LCMV Cl 13 infected WT and Sphk2-/- neutrophils. ***p≤0.001, **p≤0.01, *p≤0.05, bidirectional, unpaired Student's t-test.

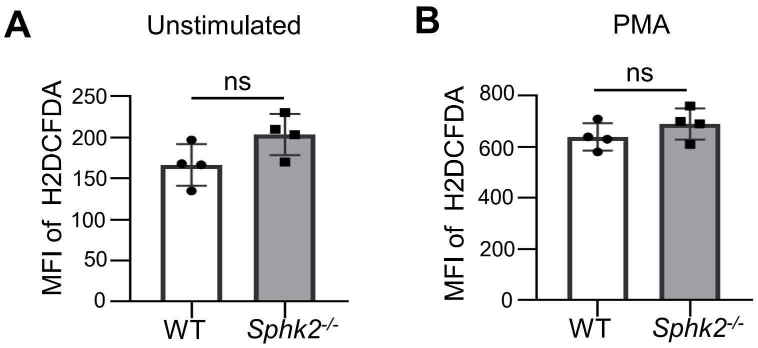


**Fig. S5. Production of ROS is not different between uninfected WT and *Sphk2$^{-/-}$* neutrophil.** Bone marrow neutrophils were isolated from uninfected C57BL/6 and *Sphk2$^{-/-}$* mice (n = 4). BMNs were cultured in the absence (**A**) or presence of 50 nM PMA for 30 min at 37°C in a CO2 incubator (**B**), and production of intracellular ROS was detected by incubating the cells with H2DCFDA for 30 min at 4°C in the dark. The mean fluorescence intensities (MFIs) of H2DCFDA were detected by flow cytometry. *n.s.* not significant, bidirectional, unpaired Student's *t*-test.

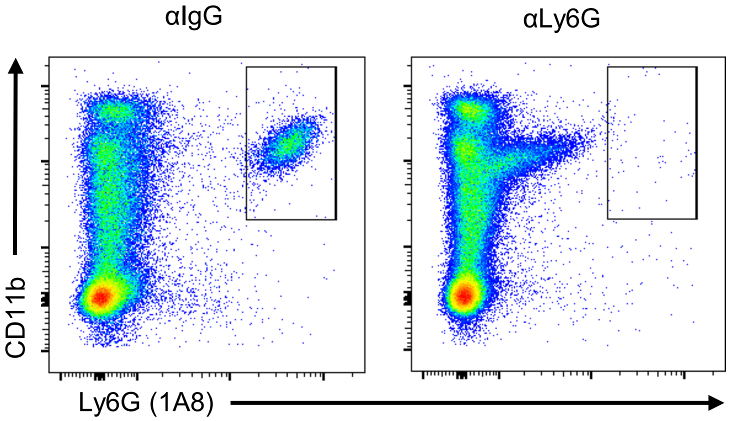


**Fig. S6. Anti-Ly6G antibody treatment depletes the neutrophils.** WT mice (n = 6/group) were infected with LCMV Cl 13 and were treated with i.p. injection of mice with 250 µg IgG isotype control or anti-Ly6G antibody (clone 1A8). 3 days post antibody treatment, 50 µl venous blood was collected from both groups and the percentage of neutrophils ($CD11b^{+}Ly6G^{+}$ cells) was quantified by flow cytometry.

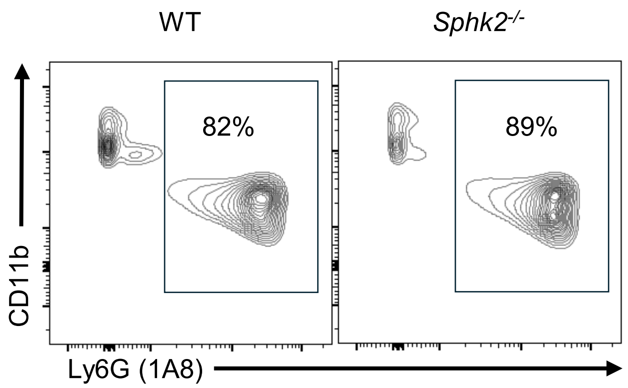


**Fig. S7. Quantifying the purity of enriched bone marrow neutrophils (BMNs).** Bone marrow neutrophils were isolated from WT and *Sphk2*$^{-/-}$ mice as described in the method section. The purities of enriched BMNs (CD11b$^+$Ly6G$^+$) were quantified by flow cytometry.

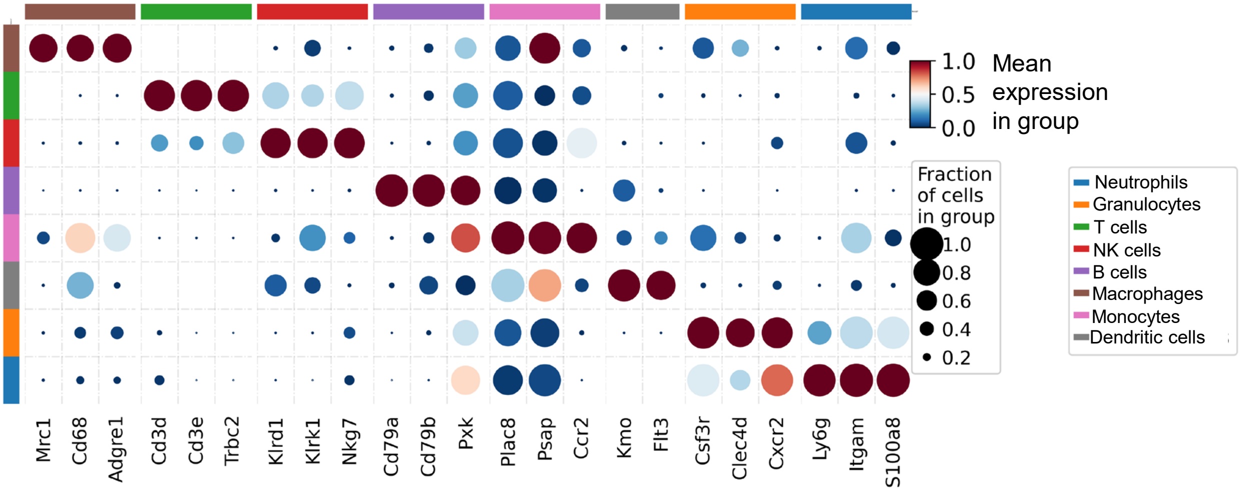


**Fig. S8. Annotation of bone marrow cell types.** Bone marrow cell types were annotated based on the expression of cell-specific marker gene or combination of gene expression profiles of specific cell subtypes.

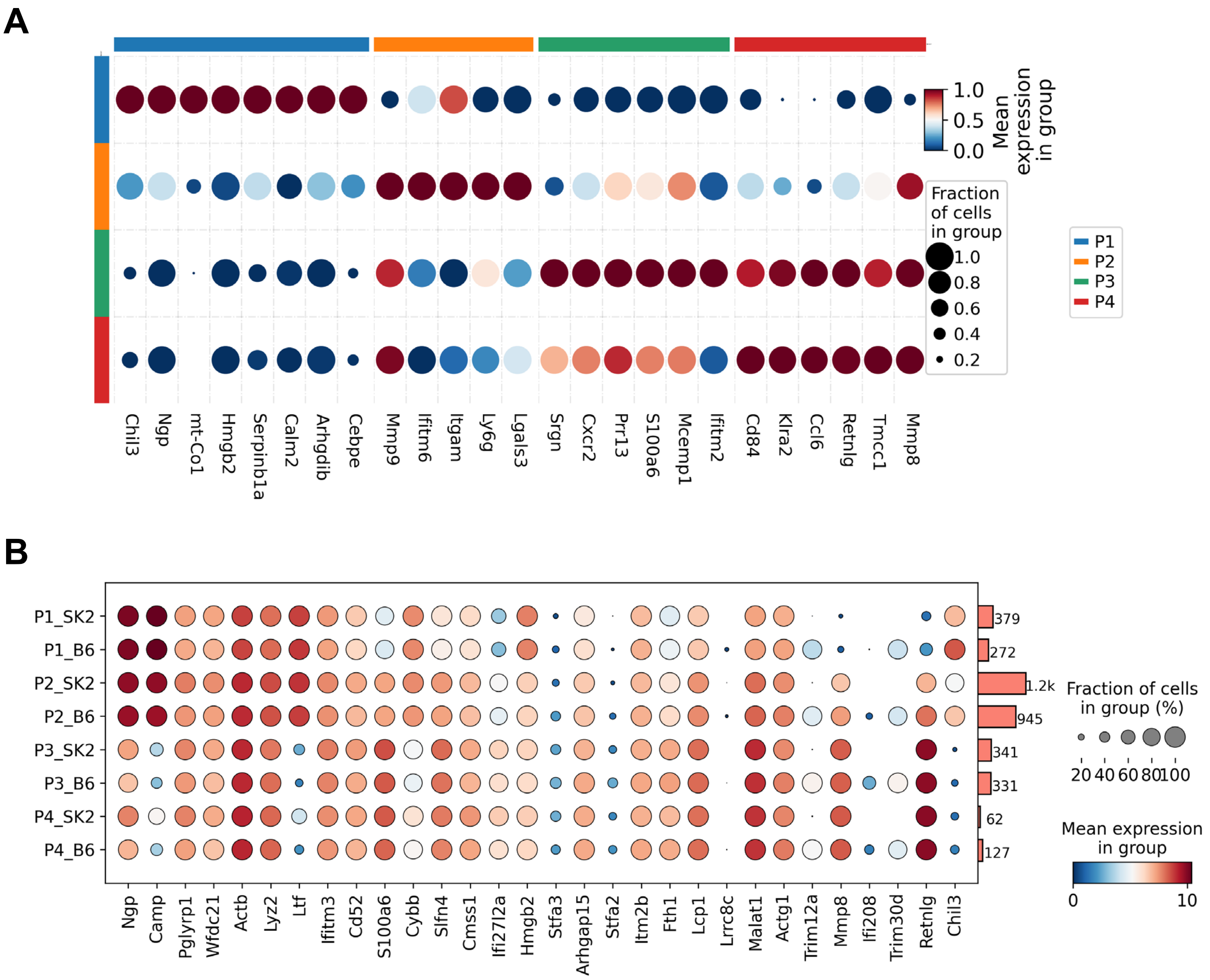


**Fig. S9. Neutrophil subtyping and distribution of top 30 DEGs on neutrophil subtypes.** Bone marrow neutrophils were classified into P1-P4 subtypes based on their gene expression profile (**A**). Comparison top 30 DEGs distribution on different subtypes of neutrophils between WT and *Sphk2*$^{-/-}$ (**B**).

**Supplementary table S1**: Antibodies used in this study.

| Antibody | Clone | Maker | Catalog |
|---|---|---|---|
| CD45-BV785 | 30-F11 | Biolgened | 103147 |
| CD11b-PerCPCy5.5 | M1/70 | Biolegend | 101230 |
| Ly6G-APC | 1A8 | Biolegend | 127614 |
| Ly6C-PE | HK1.4 | Biolegend | 128008 |
| Ly6C-BV605 | HK1.4 | Biolegend | 128036 |
| CD244-PECy7 | m2B4 | Biolegend | 133511 |
| CD244-FITC | m2B4 | Biolegend | 133503 |
| CXCR2-FITC | SA045E1 | Biolegend | 149607 |
| CD115-BV421 | AP598 | Biolegend | 135513 |
| Ly6C-FITC | HK1.4 | Biolegend | 128005 |
| CD48-PE | HM48-1 | Biolegend | 103405 |
| CX3CR1-FITC | SA011F11 | Biolegend | 149019 |
| CD107-PE | 1D4B | Biolegend | 121611 |
| CD101-PE | moushi101 | Invitrogen | 12-1011-80 |
| CD177-Alexa@647 | Y127 | BD Bioscience | 566599 |
| CD206-BV421 | C068C2 | Biolegend | 141717 |
| CD38-FITC | 521016F | Biolegend | 165608 |
| CD3-PerCPCy5.5 | 17A2 | Biolegend | 100218 |
| CD8-FITC | 5H10-1 | Biolegend | 100804 |
| CD4-FITC | RM4-5 | Biolegend | 100510 |
| CD45-FITC | S18009F | Biolegend | 157214 |
| CD33-APC | W18124D | Biolegend | 104806 |
| Ly6G-BV421 | 1A8 | Biolegend | 127627 |
| CD19-APC | 6D5 | Biolegend | 115511 |
| CD4-APC | GK1.5 | Invitrogen | 17-0041-82 |
| IFN-γ-PE | XMG1.2 | BD Bioscience | 554412 |
| CD3-PE | KT3.1.1 | Biolegend | 155607 |
| MHC-II-PE | AF6-120.1 | BD Bioscience | 553552 |
| Tim-3 | B8.2C12 | Biolegend | 134010 |
| PD-1-BV785 | 29F.1A12 | Biolegend | 135225 |
| CD8-BV605 | 53-6.7 | Biolegend | 100744 |
| TNF-α-APC | MP6-XT22 | Biolegend | 506307 |
| Granzyme B-PE | QA18A28 | Biolegend | 396405 |
| Granzyme B-APC | QA18A28 | Biolegend | 396408 |
| Ki67-FITC | 16A8 | Biolegend | 652410 |
| CD4-BV605 | RM4-5 | Biolegend | 100548 |
| TNF-α-BV605 | MP6-XT22 | Biolegend | 506329 |
| VL-4 | PS/2 | BioXcell | BE0071 |
| Rabbit anti-mouse-HRP | | Cell Signalling Technology | 7076 |
